# Clone-level multi-modal prediction of tumour drug response

**DOI:** 10.64898/2026.05.06.723206

**Authors:** Quentin Duchemin, Daniel Trejo Banos, Anne Bertolini, Pedro F. Ferreira, Rudolf Schill, Matthias Lienhard, Rebekka Wegmann, Tumor Profiler Consortium, Berend Snijder, Daniel Stekhoven, Niko Beerenwinkel, Franziska Singer, Guillaume Obozinski, Jack Kuipers

## Abstract

Tumour heterogeneity presents a major challenge for precision oncology, as genetically and phenotypically distinct tumour clones may respond differently to therapy. To address this, we introduce scClone2DR, a probabilistic multi-modal framework that predicts drug responses at the level of individual tumour clones by integrating single-cell DNA and RNA sequencing with ex-vivo drug-screening data. In simulations, scClone2DR substantially outperforms alternatives in recovering true drug-effects and clonal sensitivities. Applied to 60 melanoma and 21 acute myeloid leukaemia patient samples, the method identifies heterogeneous clonal responses, yields biologically meaningful feature rankings, highlights clones that may be resistant to treatment, and improves the prediction of clinical outcomes compared to models ignoring clonal structure. These results demonstrate that modelling tumour evolution and clonal diversity is crucial for accurate drug-response prediction and provides a foundation for more effective, clone-aware precision oncology.

## Introduction

Cancer is a complex set of diseases where diverse genomic aberrations induce characteristic hallmarks allowing for abnormal cell growth and tumour progression [1]. High-throughput methods to profile tumours have ushered in the era of precision oncology, where, despite notable successes, several hurdles remain to optimise the matching of targeted therapies to each individual patient and their tumours [2]. One strategy to match therapies to tumours is single-cell functional precision medicine (scFPM) where various treatments are tested ex-vivo on a patient’s cancer cells. This has been shown to offer clinical benefits in haematological cancers [3, 4, 5] and is currently being tested in multi-centre randomised trials [6]. Recently, scFPM has been shown to extend to drug-repurposing for brain tumours [7].

Cancer is also an evolutionary disease [8] and it has long been understood that intra-tumour heterogeneity is a serious hurdle for effective treatment and a route to relapse [9, 10]. Indeed, the realisation of precision oncology may require targeting the individual clones in a tumour [2] whose differential responses may confound scFPM while also determining long-term patient response.

A central technology to measure and resolve the genotypic diversity has been single-cell DNA (scDNA) sequencing (scDNA-seq) [11] for which approaches scale to hundreds or thousands of singlecells [12, 13]. Due to the specific noise of scDNA-seq, phylogenetic methods [14] have been specialised to the task of reconstructing tumour architectures[15, 16, 17, 18, 19, 20]. Along with genomic changes during the underlying evolutionary process, the heterogeneity of phenotypic changes can be profiled through single-cell RNA (scRNA) sequencing (scRNA-seq) [21]. In single-cell measurements, specific noise and analytical challenges arise [22, 23] in characterising the different tumour cell populations, as well as immune cells in the tumour micro-environment.

In this work, we tackle the interplay between tumour heterogeneity and drug response by integrating scFPM with clonally-resolved tumour populations obtained from scDNA and scRNA sequencing. In particular, we reconstruct the tumour phylogeny from scDNA data [14, 20] whose clones are annotated with expression and pathway information from the scRNA data [24, 25]. We embed gene and pathway features with a Variational Auto-Encoder (VAE), which is a widely adopted approach [26] for dimensionality reduction, and build on the rich history of machine learning approaches for predicting drug response trained on cell-line data [27]. Our task though runs much deeper since different individual cells are profiled with each technology, and we must first match and integrate the input data. Then we must account for the tumour heterogeneity in our prediction modelling even though we do not directly measure the responses of different populations. As such, we develop scClone2DR, a multi-modal probabilistic model for predicting drug response at the level of individual tumour clones. We retain interpretability with pathway features, uncertainty quantification, and show strong advantages over alternatives in simulation settings. Importantly, there are currently no directly comparable models for this task, as the combination of multi-modal data integration and subpopulation-level predictions is itself new. For this reason, foundation models are not yet a viable alternative, despite their recent prominence, and may only become useful once sufficient data are available for their training.

We apply our method to data collected as part of the Tumor Profiler Study [28], a large-scale multi-modal profiling of tumours within clinically-relevant timelines [29, 30, 31]. We analyse the 60 melanoma [31] and 21 acute myeloid leukemia (AML) [29] patient samples which have been profiled with scFPM, scDNA-seq and scRNA-seq (see Data description in the Methods). We demonstrate that we can resolve the differential responses of tumour sub-populations to various treatments. Not only do we find more biologically relevant response predictions, we often observe that there are tumour clones predicted to be resistant to a therapy targeting the tumour as if it were homogeneous. Moreover, by accounting for the clonal structure of each tumour we can predict the actual patient response to treatment notably better than approaches that only model average tumour features. On the AML cohort, we demonstrate how our approach can be flexibly adapted to work with tumour populations defined only with scRNA data.

## Results

### Model overview

One input to our model is the evolutionary history of each tumour reconstructed from scDNA-seq data [20] allowing us to resolve different tumour clones and their copy number profiles. The expression profiles and pathway features, obtained from the scRNA-seq data [24], are then mapped to these tumour clones [25], producing a comprehensive profile for each clone. These may be reduced to a lower dimensional representation through a VAE. The other input data are the aggregate responses of the tumour cells measured via ex-vivo drug screening [3]. Our model, scClone2DR (Methods), learns clone-specific features that explain the observed aggregate drug responses, while simultaneously modelling the drug effects that interact with these features to determine the survival of each clone under each treatment. Overall we create a probabilistic mapping from features to drug response, enabling prediction at the level of individual tumour clones (Figure 1). The output is a clone-level multi-modal prediction of response to each drug. The resulting model is further validated on the clinical response of each patient to different therapies.

**Figure 1:**
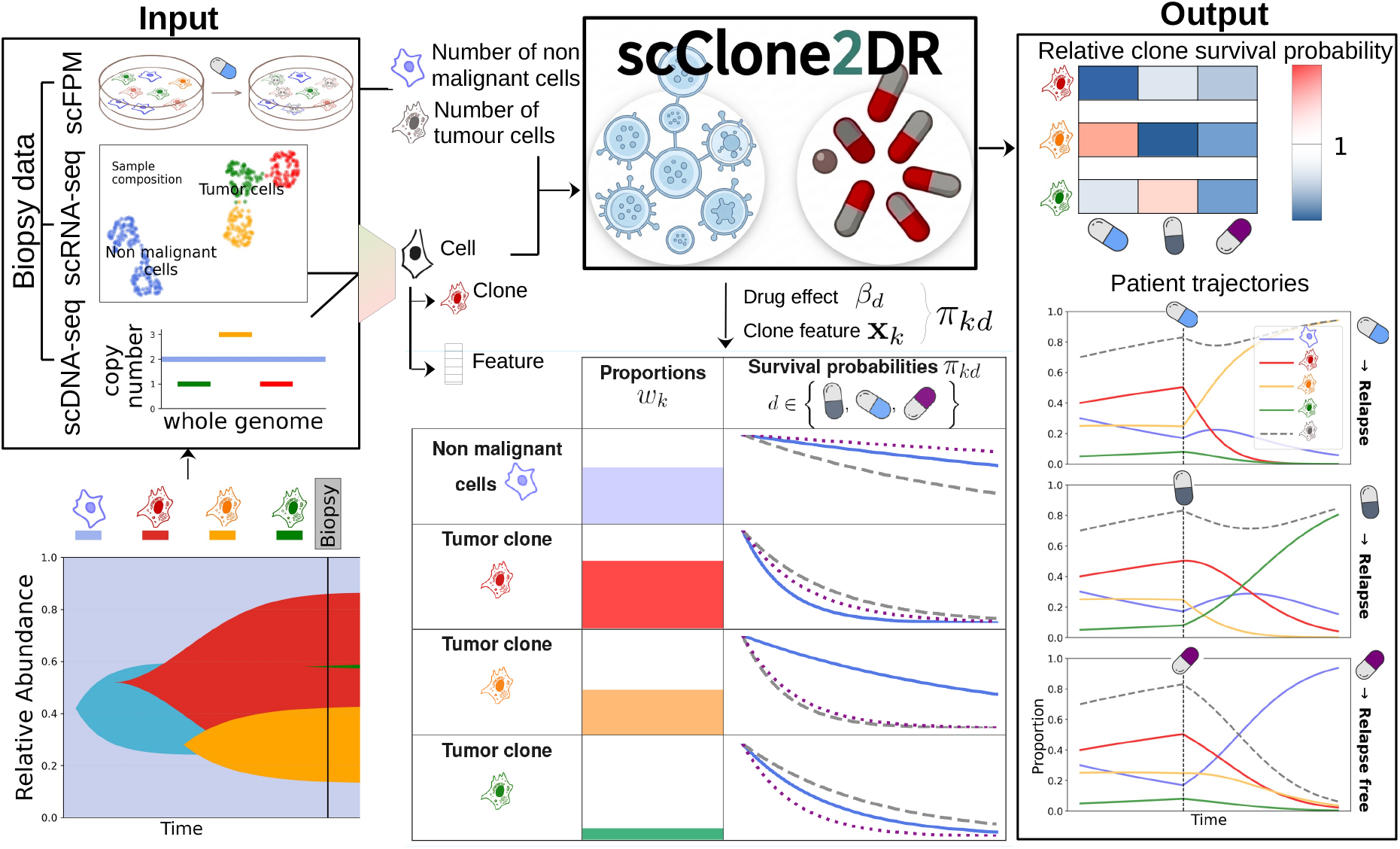
Overview of scClone2DR. The model takes scDNA-seq, scRNA-seq and scFPM drug screening as data inputs and integrates them to predict the drug response for each tumour clone. From scDNA-seq different clones are inferred from their copy number profiles which are matched to the expression profiles from scRNA-seq. Single-cells are also exposed to drugs in the ex-vivo drug screening where the survival of malignant and non-malignant cells are measured. Accounting for the gene and pathway features of the tumour clones, along with overdispersion and adjusting for confounding, scClone2DR is a comprehensive generative model for the response of each clone to each drug, extracting the effect of each drug and how this depends on the clone features. The predicted sensitivity or resistance of clones further predicts the clinical response of patients to their given treatments.

### Performance advantages in simulated settings

To evaluate the performance of scClone2DR, we ran simulation studies comparing to a range of alternatives. We provide full information on the simulation setting and alternatives in Supplementary Section S1. Mimicking values observed in the real melanoma cohort, we generate synthetic scFPM data of aggregate drug response based on the underlying simulated tumour clones and the drug-effects linked to clonal features representing pathway scores. First, we include a simple baseline which does not account for gene/pathway features and has a common drug effect across patients for malignant cells. Then we include two pseudo-bulk approaches, one (bulk) where the input features are averaged across all cells and one (dual bulk) where we have two sets of input features averaged across all malignant cells and non-malignant cells respectively. We further compare the probabilistic model of scClone2DR to machine learning approaches for the predictive task where we input the same clonal features from scClone2DR into a factorisation machine (FM) akin to the scDrug approach [32] and into a neural network (NN).

We observe much better concordance between simulated and inferred drug-effects with scClone2DR compared to the alternatives (Figure 2a, scClone2DR explains over 80% of the variance compared to less than half with any alternative; fold change is the difference between the mean log-fraction of nonmalignant cells in control and treated wells, see Supplementary Section S1). Indeed, both machine learning prediction approaches (NN and FM) recover very little of the underlying true effects, being outperformed by the simple baseline model, with the pseudo-bulk approaches performing better still.

**Figure 2:**
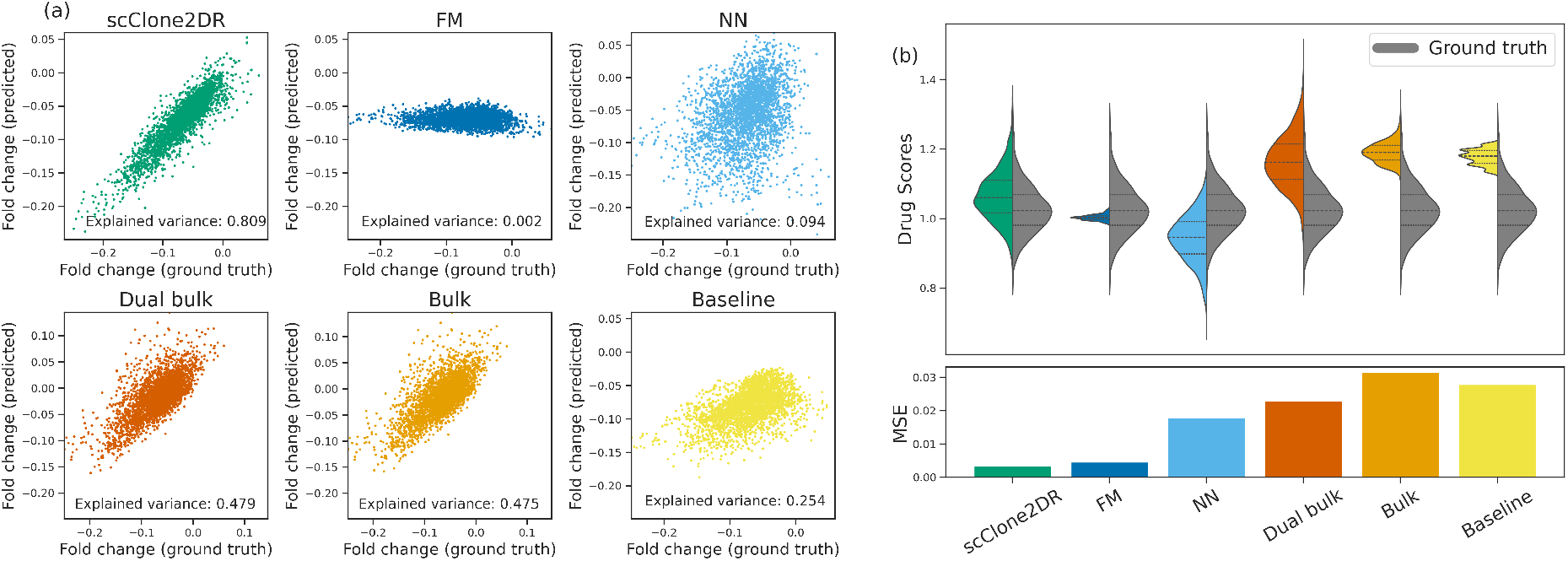
Simulation comparison. (a) Comparison of simulated and inferred drug-effects in terms of fold-changes for scClone2DR and a range of alternatives including a factorisation machine (FM) and a neural network (NN). (b) Comparison in terms of simulated and inferred drug scores, including the mean squared error (MSE).

In terms of drug scores, a measure to reflect the prediction for the most resistant tumour population (Supplementary Section S1), we again see good performance of scClone2DR (Figure 2b). Both the NN and the dual-bulk approach predict a reasonable distribution of effects, but shifted and hence with a large average error. The bulk and baseline models fail to capture the distribution while also having a large error. The FM predictions are tightly centred, allowing reasonably low typical errors, but the approach does not make relevant predictions as also observed for the fold changes (Figure 2a).

When simulating data with additional replicates of the treatments and with less overdispersion, we observe similar results (Supplementary Figure S1). While the average drug-score errors are slightly reduced for all methods, only scClone2DR is able to fully leverage the higher quality data to achieve better correlation between the inferred and simulated parameters. Further comparison metrics are presented in Supplementary Section S1, while consistency, robustness and an ablation study of scClone2DR are presented in Supplementary Section S2.

### Melanoma cohort analyses

We applied scClone2DR to the Tumor Profiler melanoma cohort of 60 samples profiled with scDNA-seq, scRNA-seq and scFPM.

#### scClone2DR for personalised cancer treatment

The main output of scClone2DR is the relative survival probabilities of different tumour clones for each drug for a given patient sample. This allows us to prioritise different treatments, for example by ranking by the weighted average effectiveness across all tumour clones (Figure 3). Compared to pure scFPM screening, we now account for the noise and make predictions for each clone. For example, the best drug for predicted aggregate effect (Elesclomol, Figure 3) has two tumour clones which are predicted to be resistant to that treatment. These may then be of concern for clinical response, and preference may be given to drugs (or combinations) to which all clones might be sensitive even if the average effects are weaker (like the third drug in Figure 3).

**Figure 3:**
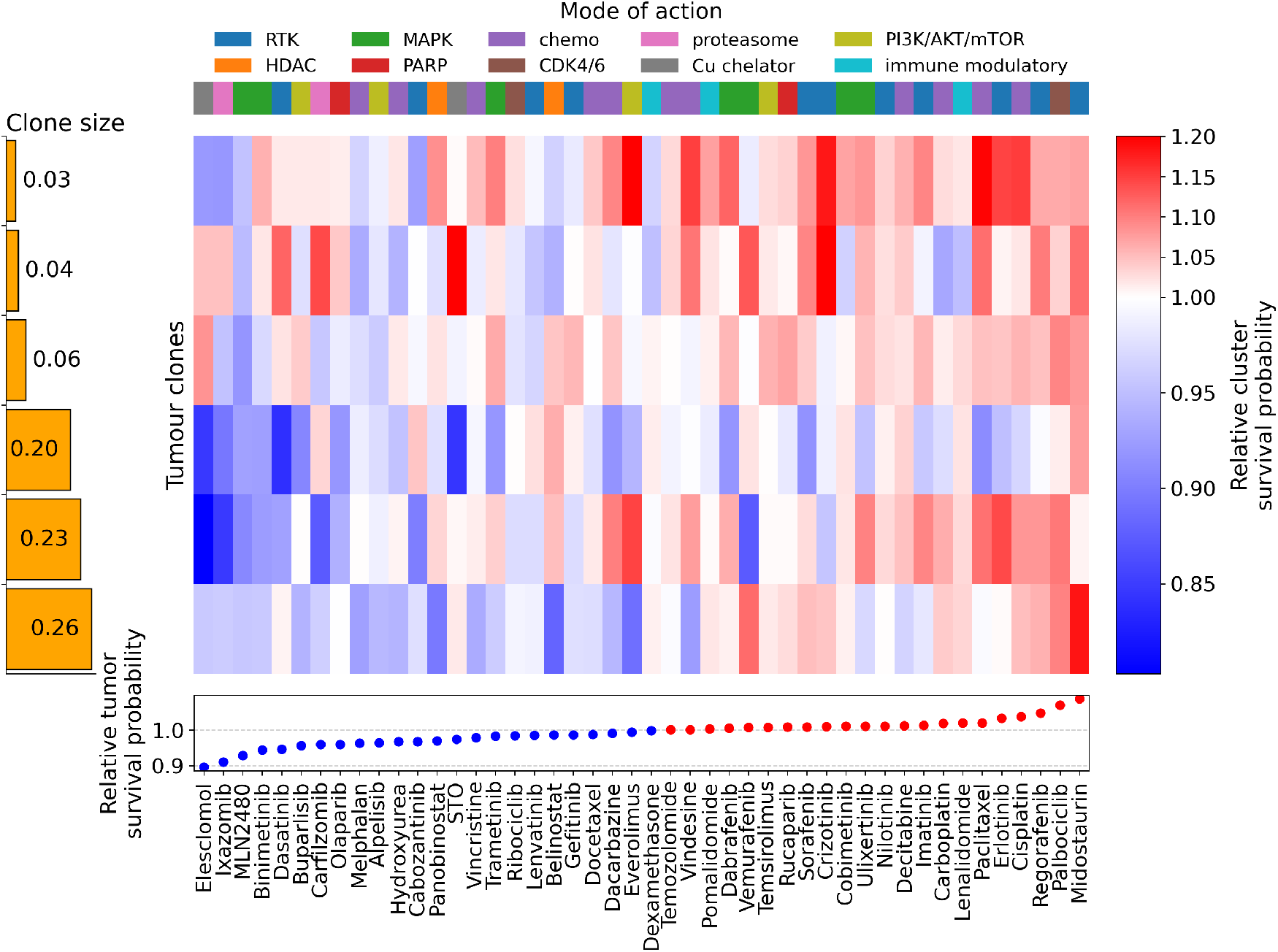
scClone2DR result for a melanoma patient. Predicted survival probabilities of each tumour clone (relative to non-malignant cells) across the range of drugs tested for patient sample MOTAMUH, with red indicating resistance and blue sensitivity in the heatmap. Drugs are ranked by weighted averages over all tumour clones.

#### Integrating scDNA and scRNA data improves performance

As a first test of performance on the real data, we explored whether using tumour clones defined through integrating scDNA and scRNA analyses (Methods) outperformed clones defined using scRNA-seq clusters. Over 100 repetitions with 60% training data (with replacement) we compute the correlation on the 40% of held-out patients between the observed fraction of tumour cells for different patients and drugs and the predicted values output by scClone2DR. With the integrative analysis the correlations were over 0.9 (Supplementary Figure S2a) and significantly better than only using scRNA-seq data (*p*-value = 5.69 × 10^−5^, Wilcoxon two-sided signed-rank test) with a clear improvement 65% of times. This indicates that the tumour clones anchored by the evolutionary history reconstructed from scDNA-seq are more relevant for predicting clonal response.

#### Pathway features retain performance and interpretability over gene-based features

With the same training-test splits we additionally compared the performance of scClone2DR when we provide clone features at a pathway level or at a gene level (Methods). Correlations tend to decrease notably when using the gene-based features with the pathway-based model routinely being as good or better (Supplementary Figure S2b), and significantly better overall (*p*-value = 1.87 × 10^−8^, Wilcoxon two-sided signed-rank test). The gene level features are still high-dimensional, so we reduce their dimensionality with a VAE (Methods). In the low-dimensional representation, we observe very similar performance to the pathway-based model with high correlation (Supplementary Figure S2c). There is a typically small improvement in performance, which is significant (*p*-value = 6.19 × 10^−5^, Wilcoxon two-sided signed-rank test), but given the small effect we prefer to retain the interpretability of the pathway-based features.

#### scClone2DR finds more biologically meaningful drug predictions

Along with the relative survival probabilities of the different tumour clones for each drug (Figure 3), scClone2DR also provides a ranking of which features were important to the prediction. The analysis was performed at the gene-level using the gene expression profile for each clone and also of the non-malignant cells (Methods). We used these gene features to assess how biologically meaningful scClone2DR results are in comparison to the dual bulk model with a single tumour clone compared to the non-malignant one. For each clone we find the best drug, either for every clone with scClone2DR or for the tumour as a whole for the bulk comparison. For each drug we retrieved a set of clinically relevant genes from the curated CIViC database [33] and evaluated if the model correctly identifies genes among this set as important for survival by computing its normalised enrichment score (NES) (Methods).

The predictions at the clone level outperform treating the tumour as whole (Figure 4a) with a significant improvement of scClone2DR over the dual bulk model of enrichment scores for the drug-specific gene sets (*p*-value = 0.0016, Wilcoxon two-sided signed-rank test). Along with the typical improvement on average, we observe more pronounced improvements for certain drugs (like Everolimus, Figure 4b). Supplementary Table S1 shows an overview of which drugs scClone2DR selects as best across all the clones of the melanoma cohort. On average, samples have between 5 and 6 tumour clones, for which scClone2DR predicts around 3 different drugs on average. This underlines the importance of respecting the heterogeneity of tumours in cancer treatment: a single drug can rarely be expected to have an effect of all tumour populations. In practice, this scenario corresponds to choosing a combination therapy to address a higher fraction of the tumour. For comparison, if we are forced to fix one drug per sample ignoring heterogeneity and targeting all tumour populations with the same treatment, we observe a subset of tumour clones where scClone2DR clearly predicts that they would not respond to the chosen treatment (Supplementary Figure S3).

**Figure 4:**
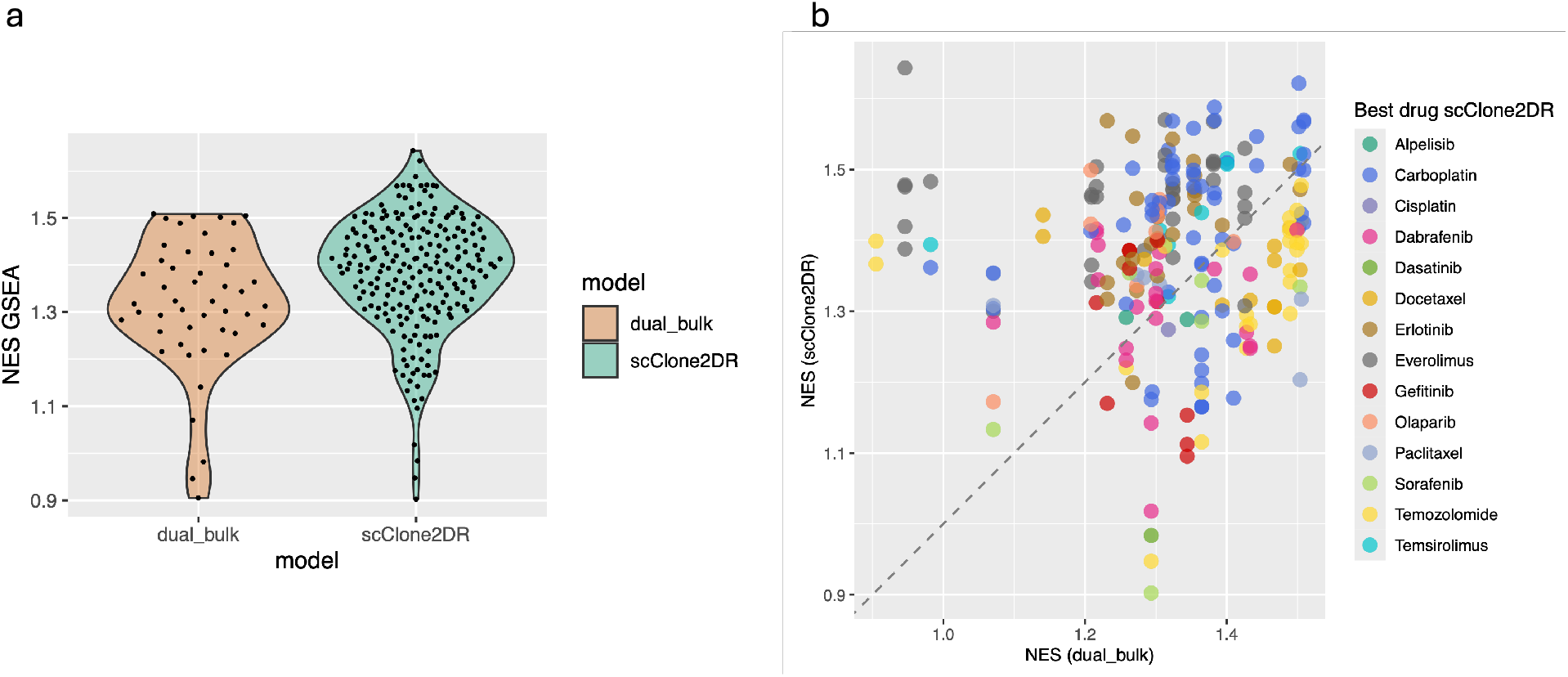
CIViC-based assessment of biological relevance. Comparison of enrichment scores of clinically relevant gene sets between scClone2DR and the dual bulk model using the gene features reported as important for drug sensitivity. The violin plot in (a) shows for the tumour clone (dual bulk) or each tumour clone (scClone2DR) the gene set enrichment score for the drug with the best score. The corresponding scatter plot in (b) additionally shows for each clone which drug was predicted as best according to scClone2DR. NES = Normalized Enrichment Score

We additionally show that inference with scClone2DR is stable, that its uncertainty quantification is well-calibrated and the importance of accounting for the experimental design in the modelling in Supplementary Section S3.

#### Clonal modelling aids predicting clinical response

As scClone2DR models drug response at the clonal level, a key question is whether the predicted sensitivity or resistance of each clone to each drug translates to the actual clinical response of patients to their therapies. From clinical response modelling (Supplementary Section S4), we observe that using the measured ex-vivo drug response values offers a slight improvement over a baseline using clinical information but still has little predictive power of actual patient outcome (Figure 5, red and yellow). Instead, accounting for the clonal structure of each tumour offers significant improvement in predictive performance (pairwise DeLong tests [34]; Supplementary Table S2). Here, the actual internal prediction engine seems to matter less (with scClone2DR a bit better than an FM; Figure 5, green and blue) while accounting for clones is essential.

**Figure 5:**
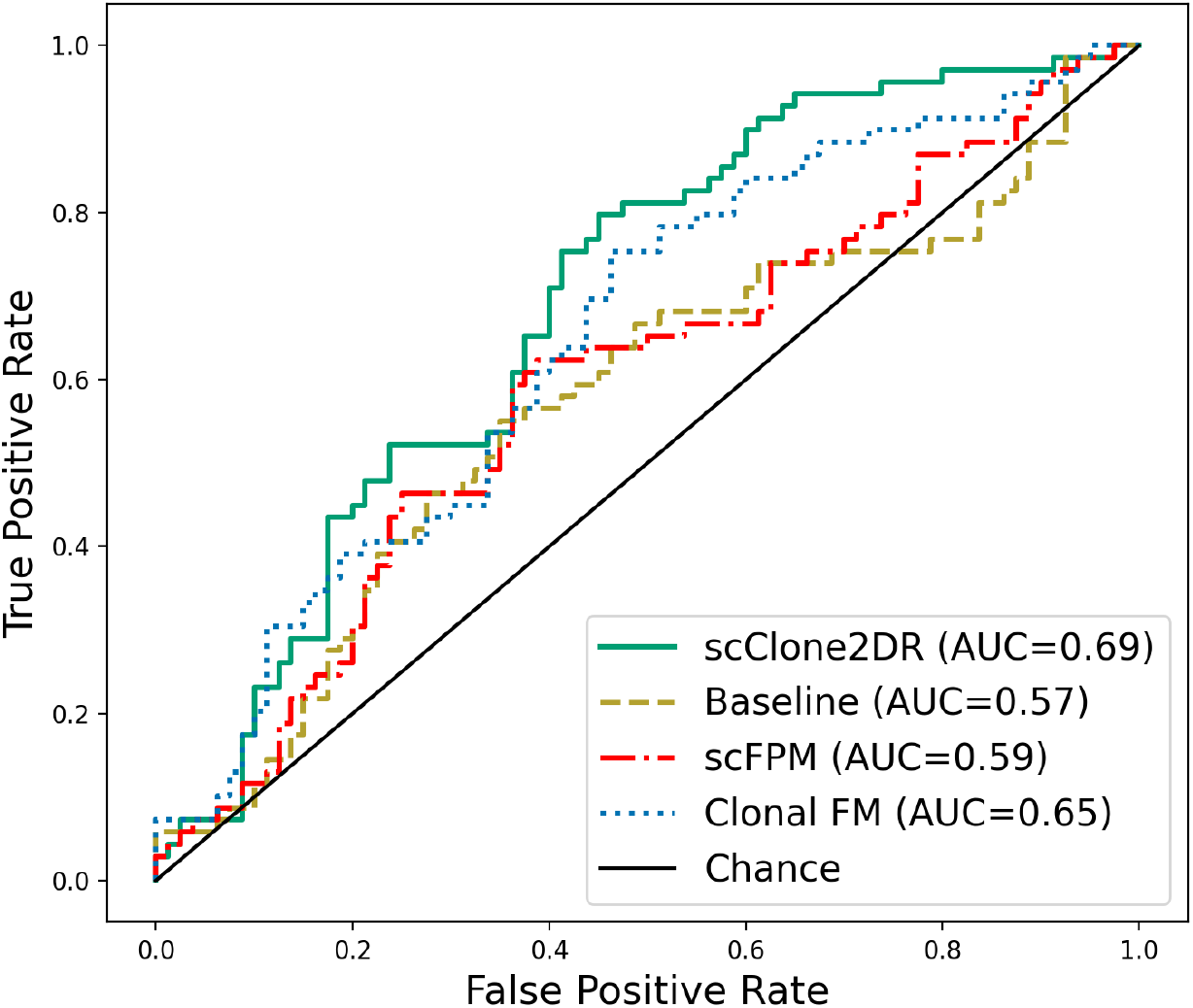
ROC curves of clinical response prediction for different models. Results are aggregated predictions across all follow-up time points and all patients for the treatments the patients actually received. This is computed using a Leave-One-Out scheme where one patient at a time is held out.

For clinical response modelling, we included two features corresponding to the predicted aggregate sensitivity/resistance of all tumour clones and that of the most resistant tumour clone. At the shortest follow-up (3 months), the aggregated feature was more relevant, while for longer times (6–9 months) the most resistant clone became of greater importance (Supplementary Figure S4). Though for longer time horizons the relevant importance keeps swapping, this may indicate that the evolutionary dynamics of tumours plays a role with resistant clones potentially expanding over longer times, a conjecture in line with the literature [8, 35].

### AML cohort analysis

We also extended scClone2DR to work for different indications with specific modelling challenges. In particular we adapted the framework to handle the AML cohort [29] by (*i*) accounting for putative cancerous cells which are classified through the cell-typing as a distinct class to the cancer and non-malignant cells, (*ii*) allowing several non-malignant clusters to account for different types of blood cells, and (*iii*) working with clusters defined from scRNA-seq data due to the sparsity of CNAs to anchor the cancer clones (Methods).

Our model can create personalised predictions for each patient for each cell type and cancer and putative cancer clones (akin to Figure 3). Similarly, we can explore the predictions per drug to see which patients exhibit sensitivity of tumour and putative tumour populations to the drug, and where the drug is predicted to be harmful for non-malignant cells. For the drug example of Venetoclax (Figure 6), we observe that a small fraction of putative cancer clones and cancer clones are predicted to be resistant (6 out of 77; 8%). Among the eleven instances where patients received the drug after sampling, five had all (putative) cancer clones with predicted relative survival below 0.9 (sensitive to the drug), only one showed a cluster with predicted relative survival above 1.1 (resistance), with the rest inbetween. In general, we also observe the heterogeneity in the predicted sensitivity of different cancer populations even within one sample.

**Figure 6:**
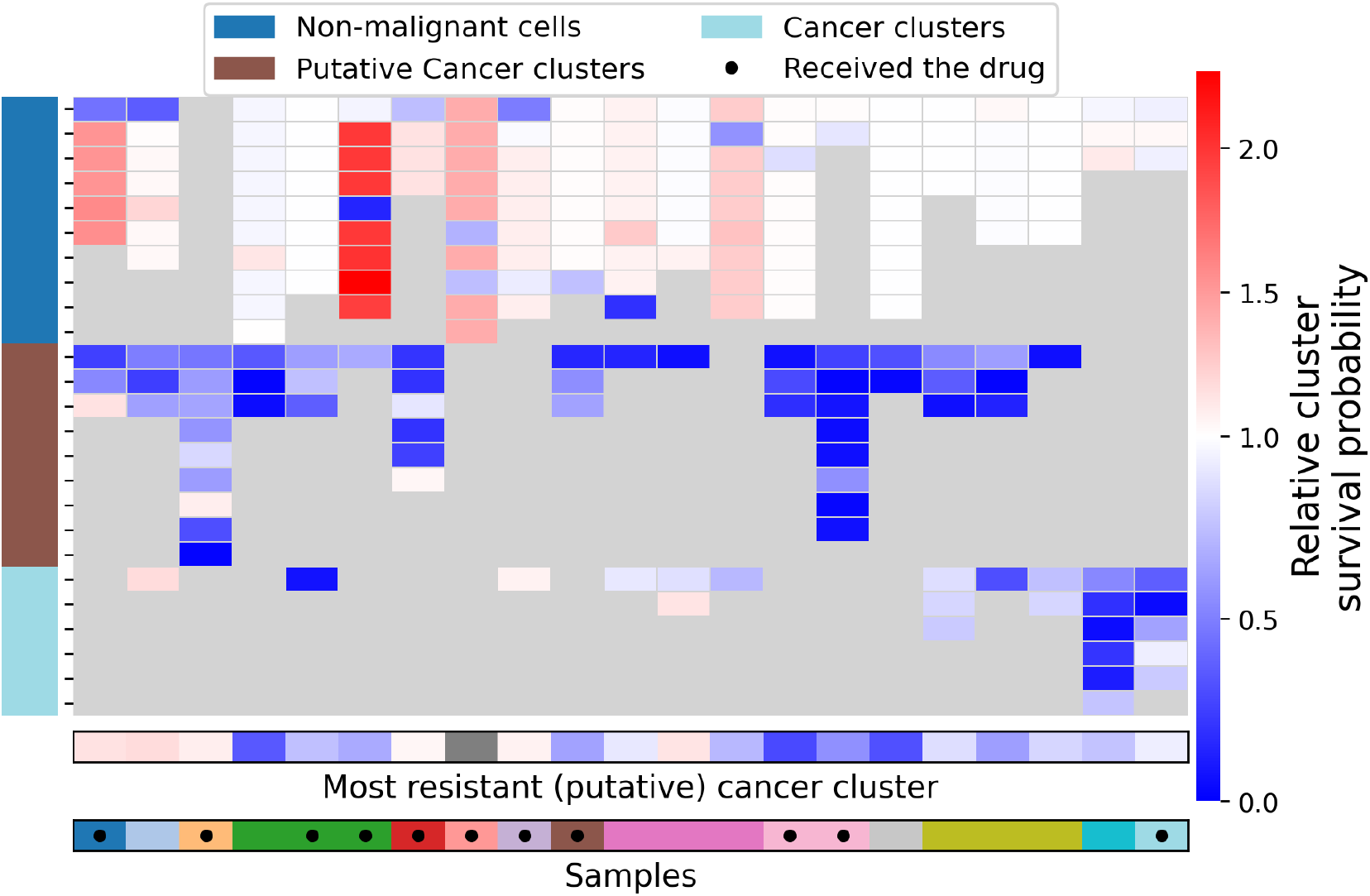
scClone2DR result for Venetoclax in AML. Relative survival probabilities for the AML cohort of cancer and non-malignant clusters in response to the drug Venetoclax. The survival probability of each cluster is normalized by the average survival probability of the non-malignant cells. The grey cells are due to the number of clusters varying across patients. We highlight the most resistant cancer or putative cancer clone in the upper horizontal annotation. Samples from the same patient acquired at different time points are indicated with the same colour in the lower horizontal annotation. The black dots indicate patients who received the drug Venetoclax just after the sample collection and data analysis.

## Discussion

Tumour heterogeneity has long been recognised as an impediment for treatment [9, 10] in that to be effective treatment needs to account for all tumour clones [2]. Here we therefore developed scClone2DR to allow us to understand scFPM measurements of ex-vivo drug response through the lens of heterogeneity. Key to the success was the integration of scDNA-seq and scRNA-seq data from multi-modal profiling of tumour samples to best define tumour clones and their expression features. The gene feature space is still high-dimensional, especially compared to the number of patient samples, necessitating a lower-dimensional projection either through pathways or methods like VAE. We found similar performance with both approaches, preferring the interpretability of pathway scores.

For the clonal prediction modelling itself we found an attention mechanism (akin to [36]) most useful, though machine learning approaches for these kinds of prediction modelling are rapidly developing [26, 27]. Thanks to the more modular design of scClone2DR, advances in embeddings and increasingly complex models with larger cohorts can be directly incorporated into our framework. Care is required since complex approaches may not always outperform simple baselines [37], so we also ensured we included such baselines in our comparisons to better gauge performance.

When training on the scFPM measurements, another key to success was to account for the experimental design in the modelling framework of scClone2DR. Indeed for the hard task of predicting unseen data, not accounting for confounding can be misleading in terms of performance assessment and in creating reliable predictions [38]. We could observe that our modelling, along with being well-calibrated, offered accurate estimation of the underlying response of each tumour clone to each therapy. This can in turn inform the experimental design. Specifically, our ablation studies demonstrated that increasing the number of replicates for each drug improves overall accuracy. In practice though, this has to be balanced against either increased scFPM cost or reducing the number of drugs tested as part of the panel design.

The main output of scClone2DR is a sensitivity/resistance score of each tumour clone to each drug for each patient. Due to the tumour heterogeneity, we often also see heterogeneous predicted responses across tumour clones, potentially raising challenges in finding a single treatment encompassing the entire tumour. Clinically, this might be addressed through combination therapies, either in parallel or sequentially, or longitudinal monitoring, for example through liquid biopsies, to check if tumour clones which were predicted to be resistant are indeed progressing and treatment needs to be adapted.

The importance of accounting for all tumour clones was highlighted by our ability to translate clone-level ex-vivo drug response predictions to the actual clinical outcome of patients. Here we tested whether our predicted response to the treatments the patients actually received correlates with their future outcome. Complicating the task is that not every treatment given was tested in the scFPM panel (only 27% of patients received a treatment after the sample collection that had been tested) limiting the scope to improve beyond baseline, although accounting for drug effects through scClone2DR also allows us to more precisely model the effects of baseline factors. Clinical response is influenced by many factors (the most relevant are included in the baseline comparison) as well as by adjuvant treatments like surgery and radiotherapy, which likewise complicate the modelling. Even with these challenges we found a significant improvement when accounting for the clonal heterogeneity of each tumour. While there is still scope for further improvements, the clone-level multi-modal drug predictions of scClone2DR offer the capacity and a clear route for their realisations.

## Methods

### Data

The data was generated from single-cell suspensions of 81 patient samples (60 metastatic melanoma and 21 AML [29]) profiled with different technologies in the Tumor Profiler Study [28]. Detailed information of the data collection and processing is available [29, 30, 31] and we recap here.

Not every sample in the full cohort was processed with each technology due to their technical requirements in terms of minimum cell viability and minimum cell numbers. We only consider those samples profiled by all three of the technologies integrated in scClone2DR. For release into ex-vivo drug testing, pathologists evaluated the diagnostic material for tissue viability and tumour cell content, requiring a minimum of 20% viable tumour cells (with exclusion of samples dominated by necrosis, haemorrhage, fibrosis, or non-tumour tissue). For release to the single-cell sequencing, at least 50,000 cells were required with a minimum viability of 50%.

For each sample passing the quality control thresholds, 50,000–200,000 single-cells were processed with both the 10x Genomics scDNA protocol to uncover copy number changes in the tumour and the 10x Genomics scRNA protocol to characterise the different cell populations.

With the scDNA-seq protocol, around 200 single-cells underwent whole-genome sequencing with around 0.5-1 million reads per cell. The barcoded reads were mapped and assigned to their individual cells. The reads were binned into 20 kb genomic regions which are corrected for GC content and mapability. The adjusted number of reads in each bin reflects the underlying copy number state of that genomic region. To identify copy number aberrations, we used SCICoNE [20] to detect potential breakpoints by pooling information related to differences in read counts across all cells and to infer the phylogenetic relationships between cells.

With the scRNA-seq protocol the transcripts of around 2,000 cells were sequenced with around 50,000 reads per cell. From the RNA profiling, samples were processed using the scAmpi pipeline [24]. This includes read mapping, quality control and filtering and normalization. The extracted features per sample are the expression profiles of each cell.

Simultaneously, scFPM profiling was performed through ex-vivo drug response screening with Pharmacoscopy [3]. Different cells from the same cell suspension were incubated overnight in 384-well plates, in the presence of one of the drugs or in control wells. The number of drugs tested per sample varied as the plate design evolved over the study for melanoma, and was limited to smaller panels for samples with not enough cells in the suspension. We consider the 46 drugs that were commonly profiled in the melanoma cohort, and 69 drugs for AML, each with up to three replicates at each of two concentrations.

Automated confocal microscopy and single-cell image analysis were utilised to measure the response of the cell population at the end of the incubation period. The extracted features per sample are the total final number of tumour and non-tumour cells in wells (with or without drug compounds). Pertinent for our modelling is that measurements, being destructive, are only performed at the end, meaning that no information is available regarding the initial cell population before incubation. In particular, the total number of cells initially placed in the wells and the precise proportion of tumour cells were not measured.

### The scClone2DR approach

scClone2DR predicts drug response at the clonal level (Supplementary Figure S7 shows a technical overview), accounting for tumour heterogeneity and various noise in the data.

### Tumour clones

To learn the tumour clones we reconstruct the evolutionary history of the tumour from the scDNA-seq data [20] as performed during the Tumor Profiler analysis of each sample. This method identifies distinct clonal populations by modelling the acquisition of genomic mutations directly along the branches of the reconstructed tumour phylogeny. The processed scRNA-seq data [24] is condensed into metacells with SEACells [39] which are mapped onto the tumour clones [25].

### RNA features

The dimensionality of the metacell RNA data is first reduced by either considering pathways or genes in one of three ways:

1. We computed GSVA (Gene Set Variation Analysis) scores [40] considering different set of pathways. Unlike gene-based differential expression analysis, GSVA assesses the collective behaviour of predefined gene sets or pathways. It computes enrichment scores for each sample, quantifying how coordinated the expression of genes within a set is relative to other samples. Throughout this work, we use the hallmark gene set collection of the MSigDB resource as the source set of pathways [41, 42], which comprises 50 gene sets representing well-defined biological processes.
2. To retain the gene-level interpretability, from the normalized data from sctransform [43], we use the 825 genes from the OncoKB™ Cancer Gene List [44] which 825 were expressed in at least 95% of samples in our data set.
3. To further reduce dimensionality, we train a VAE using scVI [45] on the raw RNA metacell counts from the 825 genes mentioned above. The resulting 10-dimensional latent embedding is then used as the feature set for prediction.

The following steps of scClone2DR can proceed with any of these input RNA features.

### Clone features

Next, instead of directly using averaged (embedded) features, we keep track of the biological variability by using an attention mechanism, inspired by [36]. The attention mechanism dynamically assigns weights to cells based on their relevance, enhancing interpretability, handling heterogeneity, and reflecting contextual relationships. We present the details in Supplementary Section S5.

### Modelling

The inputs are single-cell features, with each cell assigned to either the non-malignant population or a tumour clone based on integrated scDNA and scRNA analyses. We detail the noise and graphical modelling in Supplementary Section S6 and focus here on the core of scClone2DR which is that the drug-specific survival probability for cells from sample *i* and clone *k* is modelled as

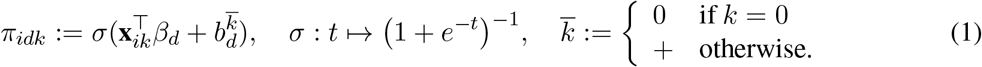

depending on the clonal features. As such we employ a generalised linear model (with a logistic link function) between clone features, **x**, and cell survival related to the drug effects, *β*, which we infer. We consider different offsets, *b*, for the drug-specific survival probabilities for the clone associated with non-malignant cells (indexed by 0) and any clones of tumour cells (indexed by *k* ≥ 1). Our model then encompasses a simple baseline model where we remove any drug-specific survival probabilities by setting *β*_*d*_ = **0** so the results are independent of the feature vector.

The final factor we consider is the dependence of the data on the experimental design of the scFPM platform, which we detail in Supplementary Section S7.

In the data, the number of replicates of control wells and treated wells for a given drug can slightly vary across patient and/or drug. Above we considered *R*_*C*_ and *R*_*T*_ independent of patient and drug to simplify notation, and we account for any variations in our code and inference. Table 1 provides the typical values of the parameters for the Melanoma cohort.

**Table 1:** Typical parameter values for the Melanoma cohort analysed.

| # replicates for control wells | # replicates for treated wells | # drugs tested | Maximum number of clones |
| --- | --- | --- | --- |
| $R_C = 24$ | $R_T = 4$ | $D = 46$ | $\max_i K_i = 9$ |

### CIViC-based evaluation

On real data, we assess the model performance in aligning with clinical knowledge of drug sensitivity or resistance based on known drug targets. We compare scClone2DR and the dual bulk model in terms of their ability to predict relevant features, as reported in the curated CIViC database [33]. Per drug we extracted the set of genes reported as relevant for that drug in at least one CIViC evidence item (regardless of disease type) to generate a CIViC-based gene set denoting the genes considered relevant for the drug response based on curated clinical evidence. For the selection of genes to consider, we used the OncoKB™ Cancer Gene List [44], of which 825 were expressed in at least 95% of samples in our data set. These were used throughout the CIViC-based analyses.

### Log odds ratio of tumour clones, relative to the non-malignant population

For the assessment of the biological relevance, we consider normalized RNA counts as features. This way, each entry of the feature vector corresponds to a single gene. To define a comparison metric we start with the odds ratio, as in [46], which tells us if it is more or less likely for cells with a specific feature to survive compared to cells without that feature. For a given clone *k* 0, the log odds ratio (LOR) for feature *j* is defined by:

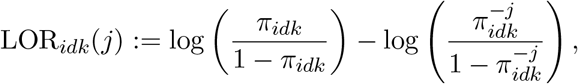

Where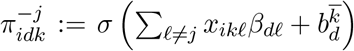 Here, 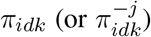 is the probability of survival for clone *k* when feature *j* is included (or excluded). An odds ratio greater than 1 (LOR *>* 0) indicates that the cells in clone *k* are more likely to survive in the presence of feature *j*. Looking at the tumour clone *k* alone might not provide the full picture since we should compare a clone *k* ≥1 of tumour cells with a reference group, namely the one of the non-malignant cells. As such we define

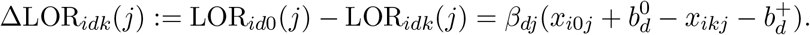

Here, LOR_*id*0_(*j*) is the log odds ratio for the group of non-malignant cells. ΔLOR_*idk*_(*j*) quantifies the extent to which feature *j* contributes to the sensitivity of clone *k* to drug *d* compared to the baseline group of non-malignant cells. If ΔLOR_*idk*_(*j*) *>* 0, it means that feature *j* contributes to increasing the sensitivity to drug *d* in tumour clone *k* compared to the reference group of non-malignant cells. For example, if using feature *j* halves the chance of survival for tumour cells in clone *k*, ΔLOR_*idk*_(*j*) will be positive only if feature *j* does not reduce the chance of survival for non-malignant cells by more than half.

- If ΔLOR_*idk*_(*j*) *>* 0, then feature *j* reduces survival in tumour cells more than in non-malignant cells (i.e., tumour-specific sensitivity).
- If ΔLOR_*idk*_(*j*) *<* 0, then feature *j* affects non-malignant cells more, which might indicate toxicity or lack of tumour specificity.

### Gene set enrichment

To assess the model performance of scClone2DR and the dual bulk model, for each drug and clone combination we performed a gene set enrichment analysis (GSEA) [42] with the FGSEA implementation [v. 1.34.0] [47] using the predicted ΔLOR per gene feature. Each vector of ΔLOR values was ranked by absolute values in descending order and with this the Normalized Enrichment Score (NES) was calculated for each CIViC-based gene set with five or more overlapping genes in our data set. Comparisons between the NES of scClone2DR and the dual bulk model were done using a two-sided Wilcoxon rank sum test.

### Model adaptations for AML

AMLs are not tumours so we refer to cancer cells and cancer clones instead (compared to the notation of Supplementary Table S3). In the AML cohort [29], copy number aberrations are less common genetic drivers of cancer progression, indeed almost half of the samples had no detected CNA. The scDNA-based analysis may then only resolve part of the heterogeneity, and we therefore focus on cancer clones as defined by clustering the scRNA-seq data using PhenoGraph [48].

Additionally, in the development of the cancer from hematopoietic stem cells there may be preleukemic cells present. In the cell-typing we therefore define clone labels: *i*) non-malignant, *ii*) putative cancer, and *iii*) cancer. Each group of cells is assigned one of these labels, and scClone2DR is modified to learn label-specific attention mechanisms and offsets to compute the survival probability for each group. The survival probability of the *k*-th group is now given by:

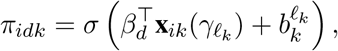

where **x**_*ik*_(*γ*_*ℓk*_) is the feature vector for the *k*-th group of cells using the parameter *γ*_*ℓk*_ in the attention mechanism,

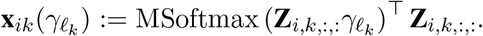

Finally, we no longer consider the non-malignant cells to be one group but now allow several clusters of cells to have the non-malignant label. This allows scClone2DR to explicitly model the different behaviour of different blood cell types in culture. The AML workflow then consists of clustering the metacells of patient *i* into clusters {0,…, *K*_*i*_} using the PhenoGraph algorithm. Labels are then assigned to each cluster, based on the dominant cell-type of each cluster to arrive at a hierarchical structure (Supplementary Figure S5). For example we obtain the clones and labels depicted in Supplementary Figure S6.

From these modifications we return to the standard scClone2DR model by reducing to only two (non-malignant or cancer) labels and having a single non-malignant population.

## Data availability

To comply with applicable laws and regulations (the Swiss Human Research Act), all de-identified data relevant to this publication will be provided as supporting information to the paper. The public AML data from [29], processed for input into scClone2DR, is available on Zenodo: https://doi.org/10.5281/zenodo.20035241.

Access to the patient-level clinical and raw biological data will be granted to registered users listed on the data access agreement with the Tumor Profiler Center (TPC) within 4 weeks of receipt of the Data Access Agreement, provided that the applicant submits all necessary ethics committee approval and supporting documents needed to meet the requirements of the agreement. Data access can be requested by contacting the TPC. The user institution agrees to destroy or discard the data once it is no longer used for the project, and in cases where data must be archived, it must be deleted within 10 years of the project’s completion. If data has not been archived, it must be deleted no later than 2 years following the completion of the project. An extension to this period can be provided upon request to the TPC leadership. Data sharing is subject to honouring patient privacy and data integrity.

## Code availability

The scClone2DR package is available on GitHub: https://github.com/cbg-ethz/scClone2DR. The package comes with a detailed tutorial that guides users through its functionality by generating simulated datasets that closely mimic single-cell data from the Tumor Profiler Study including the scFPM data. A tutorial is also provided to run the model on the AML data, using a ready-to-use version of the dataset for scClone2DR, available on the following Zenodo archive: https://doi.org/10.5281/zenodo.20035241. Within the same repository, the reference data and customized scripts used for the CIViC-based assessment of biological relevance are available.

## Acknowledgements

This project was partially funded by PHRT and SDSC (grant numbers: 2021-802 and C21-19P).

## Author contributions

Conceptualization: JK, NB, DS, BS; Data curation: AB, PFF, RW, ML, FS; Formal analysis: QD, DTB, AB; Methodology: QD, DTB, AB, PFF, RS, ML, RW, FS, GO, JK; Software: QD; Supervision: BS, NB, FS, GO, JK; Visualization: QD, DTB, AB, FS, GO, JK; Writing – original draft: QD, DTB, AB, FS, JK; Writing – review & editing: all authors.

### Tumor Profiler Consortium

Rudolf Aebersold^5^, Melike Ak^34^, Faisal S Al-Quaddoomi^12,23^, Silvana I Albert^10^, Jonas Albinus^10^, Ilaria Alborelli^30^, Sonali Andani^9,23,32,37^, Per-Olof Attinger^14^, Marina Bacac^22^, Monica-Andreea Baciu-Drăgan^7^, Daniel Baumhoer^30^, Beatrice Beck-Schimmer^45^, Niko Beerenwinkel^7,23^, Christian Beisel^7^, Lara Bernasconi^33^, Anne Bertolini^12,23^, Bernd Bodenmiller^11,41^, Ximena Bonilla^9^, Lars Bosshard^12,23^, Byron Calgua^30^, Ruben Casanova^41^, Stéphane Chevrier^41^, Natalia Chicherova^12,23^, Ricardo Coelho^24^, Maya D’Costa^14^, Esther Danenberg^43^, Natalie R Davidson^9^, Reinhard Dummer^34^, Stefanie Engler^41^, Martin Erkens^20^, Katja Eschbach^7^, Cinzia Esposito^43^, AndréFedier^24^, Pedro F Ferreira^7^, Joanna Ficek-Pascual^1,9,17,23,32^, Anja L Frei^37^, Bruno Frey^19^, Sandra Goetze^10^, Linda Grob^12,23^, Gabriele Gut^43^, Detlef Günther^8^, Pirmin Haeuptle^3^, Viola Heinzelmann-Schwarz^24,29^, Sylvia Herter^22^, Rene Holtackers^43^, Tamara Huesser^22^, Alexander Immer^9,18^, Anja Irmisch^34^, Francis Jacob^24^, Andrea Jacobs^41^, Tim M Jaeger^14^, Alva R James^9,23,32^, Philip M Jermann^30^, André Kahles^9,23,32^, Abdullah Kahraman^15,23,37^, Viktor H Koelzer^30,37,42^, Werner Kuebler^31^, Jack Kuipers^7,23^, Christian P Kunze^28^, Christian Kurzeder^27^, Kjong-Van Lehmann^2,4,9,16^, Mitchell Levesque^34^, Ulrike Lischetti^24^, Flavio C Lombardo^24^, Sebastian Lugert^14^, Gerd Maass^19^, Markus G Manz^36^, Philipp Markolin^9^, Martin Mehnert^10^, Julien Mena^5^, Julian M Metzler^35^, Nicola Miglino^36,42^, Emanuela S Milani^10^, Holger Moch^37^, Simone Muenst^30^, Riccardo Murri^44^, Charlotte KY Ng^30,40^, Stefan Nicolet^30^, Marta Nowak^37^, Monica Nunez Lopéz^24^, Patrick GA Pedrioli^6^, Lucas Pelkmans^43^, Salvatore Piscuoglio^24,30^, Michael Prummer^12,23^, Laurie Prélot^9,23,32^, Natalie Rimmer^24^, Mathilde Ritter^24^, Christian Rommel^20^, María L Rosano-González^12,23^, Gunnar Rätsch^1,6,9,23,32^, Natascha Santacroce^7^, Jacobo Sarabia del Castillo^43^, Ramona Schlenker^21^, Petra C Schwalie^20^, Severin Schwan^14^, Tobias Schär^7^, Gabriela Senti^33^, Wenguang Shao^10^, Franziska Singer^12,23^, Sujana Sivapatham^41^, Berend Snijder^5,23^, Bettina Sobottka^37^, Vipin T Sreedharan^12,23^, Stefan Stark^9,23,32^, Daniel J Stekhoven^12,23^, Tanmay Tanna^7,9^, Alexandre PA Theocharides^36^, Tinu M Thomas^9,23,32^, Markus Tolnay^30^, Vinko Tosevski^22^, Nora C Toussaint^13^, Mustafa A Tuncel^7,23^, Marina Tusup^34^, Audrey Van Drogen^10^, Marcus Vetter^26^, Tatjana Vlajnic^30^, Sandra Weber^33^, Walter P Weber^25^, Rebekka Wegmann^5^, Michael Weller^39^, Fabian Wendt^10^, Norbert Wey^37^, Andreas Wicki^36,42^, Mattheus HE Wildschut^5,36^, Bernd Wollscheid^10^, Shuqing Yu^12,23^, Johanna Ziegler^34^, Marc Zimmermann^9^, Martin Zoche^37^, Gregor Zuend^38^

^1^ AI Center at ETH Zurich, Andreasstrasse 5, 8092 Zurich, Switzerland. ^2^ Cancer Research Center Cologne-Essen, University Hospital Cologne, Cologne, Germany. ^3^ Cantonal Hospital Baselland, Medical University Clinic, Rheinstrasse 26, 4410 Liestal, Switzerland. ^4^ Center for Integrated Oncology Aachen (CIO-A), Aachen, Germany. ^5^ ETH Zurich, Department of Biology, Institute of Molecular Systems Biology, Otto-Stern-Weg 3, 8093 Zurich, Switzerland. ^6^ ETH Zurich, Department of Biology, Wolfgang-Pauli-Strasse 27, 8093 Zurich, Switzerland. ^7^ ETH Zurich, Department of Biosystems Science and Engineering, Schanzenstrasse 44, 4056 Basel, Switzerland. ^8^ ETH Zurich, Department of Chemistry and Applied Biosciences, Vladimir-Prelog-Weg 1-5/10, 8093 Zurich, Switzerland. ^9^ ETH Zurich, Department of Computer Science, Institute of Machine Learning, Universitätstrasse 6, 8092 Zurich, Switzerland. ^10^ ETH Zurich, Department of Health Sciences and Technology, Otto-Stern-Weg 3, 8093 Zurich, Switzerland. ^11^ ETH Zurich, Institute of Molecular Health Sciences, Otto-Stern-Weg 7, 8093 Zurich, Switzerland. ^12^ ETH Zurich, NEXUS Personalized Health Technologies, Wagistrasse 18, 8952 Zurich, Switzerland. ^13^ ETH Zurich, Swiss Data Science Center, Wasserwerkstrasse 10, 8092 Zurich, Switzerland. ^14^ F. Hoffmann-La Roche Ltd, Grenzacherstrasse 124, 4070 Basel, Switzerland. ^15^ FHNW, School of Life Sciences, Institute of Chemistry and Bioanalytics, Muttenz, Switzerland. ^16^ Joint Research Center Computational Biomedicine, University Hospital RWTH Aachen, Aachen, Germany. ^17^ Life Science Zurich Graduate School, Biomedicine PhD Program, Winterthurerstrasse 190, 8057 Zurich, Switzerland. ^18^ Max Planck ETH Center for Learning Systems. ^19^ Roche Diagnostics GmbH, Nonnenwald 2, 82377 Penzberg, Germany. ^20^ Roche Pharmaceutical Research and Early Development, Roche Innovation Center Basel, Grenzacherstrasse 124, 4070 Basel, Switzerland. ^21^ Roche Pharmaceutical Research and Early Development, Roche Innovation Center Munich, Roche Diagnostics GmbH, Nonnenwald 2, 82377 Penzberg, Germany. ^22^ Roche Pharmaceutical Research and Early Development, Roche Innovation Center Zurich, Wagistrasse 10, 8952 Schlieren, Switzerland. ^23^ SIB Swiss Institute of Bioinformatics, Lausanne, Switzerland. ^24^ University Hospital Basel and University of Basel, Department of Biomedicine, Hebelstrasse 20, 4031 Basel, Switzerland. ^25^ University Hospital Basel and University of Basel, Department of Surgery, Brustzentrum, Spitalstrasse 21, 4031 Basel, Switzerland. ^26^ University Hospital Basel, Brustzentrum & Tumorzentrum, Petersgraben 4, 4031 Basel, Switzerland. ^27^ University Hospital Basel, Brustzentrum, Spitalstrasse 21, 4031 Basel, Switzerland. ^28^ University Hospital Basel, Department of Information- and Communication Technology, Spitalstrasse 26, 4031 Basel, Switzerland. ^29^ University Hospital Basel, Gynecological Cancer Center, Spitalstrasse 21, 4031 Basel, Switzerland. ^30^ University Hospital Basel, Institute of Medical Genetics and Pathology, Schönbeinstrasse 40, 4031 Basel, Switzerland. ^31^ University Hospital Basel, Spitalstrasse 21/Petersgraben 4, 4031 Basel, Switzerland. ^32^ University Hospital Zurich, Biomedical Informatics, Schmelzbergstrasse 26, 8006 Zurich, Switzerland. ^33^ University Hospital Zurich, Clinical Trials Center, Rämistrasse 100, 8091 Zurich, Switzerland. ^34^ University Hospital Zurich, Department of Dermatology, Gloriastrasse 31, 8091 Zurich, Switzerland. ^35^ University Hospital Zurich, Department of Gynecology, Frauenklinikstrasse 10, 8091 Zurich, Switzerland. ^36^ University Hospital Zurich, Department of Medical Oncology and Hematology, Rämistrasse 100, 8091 Zurich, Switzerland. ^37^ University Hospital Zurich, Department of Pathology and Molecular Pathology, Schmelzbergstrasse 12, 8091 Zurich, Switzerland. ^38^ University Hospital Zurich, Rämistrasse 100, 8091 Zurich, Switzerland. ^39^ University Hospital and University of Zurich, Department of Neurology, Frauenklinikstrasse 26, 8091 Zurich, Switzerland. ^40^ University of Bern, Department of BioMedical Research, Murtenstrasse 35, 3008 Bern, Switzerland. ^41^ University of Zurich, Department of Quantitative Biomedicine, Winterthurerstrasse 190, 8057 Zurich, Switzerland. ^42^ University of Zurich, Faculty of Medicine, Zurich, Switzerland. ^43^ University of Zurich, Institute of Molecular Life Sciences, Winterthurerstrasse 190, 8057 Zurich, Switzerland. ^44^ University of Zurich, Services and Support for Science IT, Winterthurerstrasse 190, 8057 Zurich, Switzerland. ^45^ University of Zurich, VP Medicine, Künstlergasse 15, 8001 Zurich, Switzerland.

