## Supplementary figures, tables and material for "Clone-level multi-modal prediction of tumour drug response"

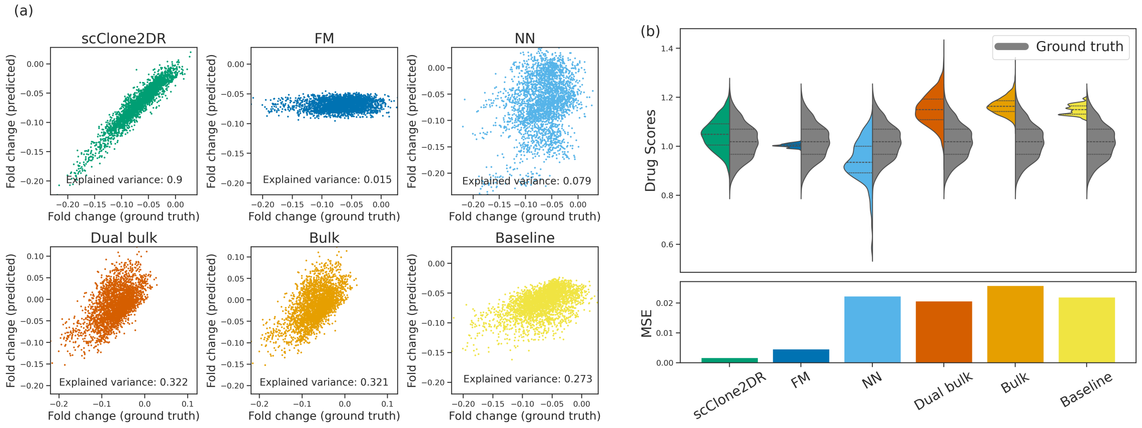


Figure S1: **Further simulation comparisons.** Counterpart of Figure 2, illustrating results with a larger number of replicates for the treated wells replicates (20 instead of 5) and a smaller overdispersion effect ($\theta = 10^5$ rather than $10^3$). (a) Comparison of simulated and inferred drug-effect parameters in terms of fold-changes for scClone2DR and a range of alternatives including a factorisation machine (FM) and a neural network (NN). (b) Comparison in terms of simulated and inferred drug scores, including the mean squared error (MSE).

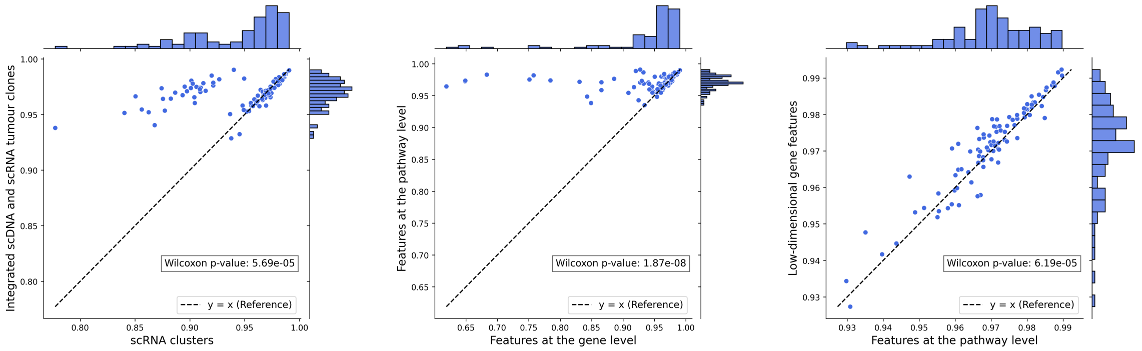


(a) (b) (c)

Figure S2: **Relative performance with different input features.** Correlation between predicted and observed tumour cell fractions on test sets. Each point represents a random selection of 60% training samples, where (a) the scClone2DR model takes as input tumour clones either defined using an integrated scDNA and scRNA analysis (vertical axis) or solely from the scRNA clusters (horizontal axis) for pathway-level features. (b) Comparing pathway-based features to gene-base features as inputs to scClone2DR (for integrated scDNA and scRNA clones). (c) Comparing pathway-based features to low-dimensional gene features obtained through a VAE (for integrated scDNA and scRNA clones).

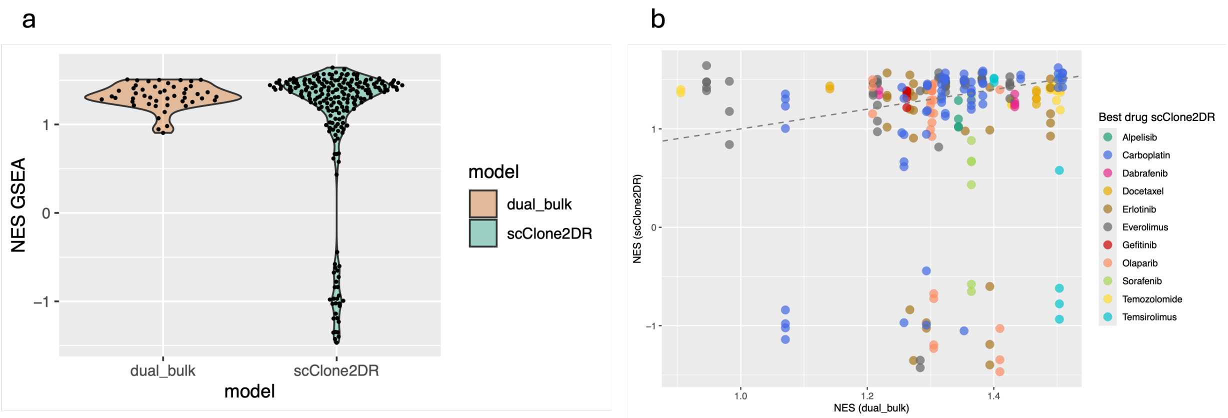


Figure S3: **CIViC-based assessment of biological relevance**. Comparison of gene set enrichment scores between scClone2DR and the dual bulk model on the gene features predicted as important for drug sensitivity. The violin plot in (a) and corresponding scatter plot in (b) shows the enrichment scores when instead of the best drug per clone only one drug is fixed across all clones of a sample, i.e., across all clones the best drug is selected for the entire sample. NES = Normalized Enrichment Score

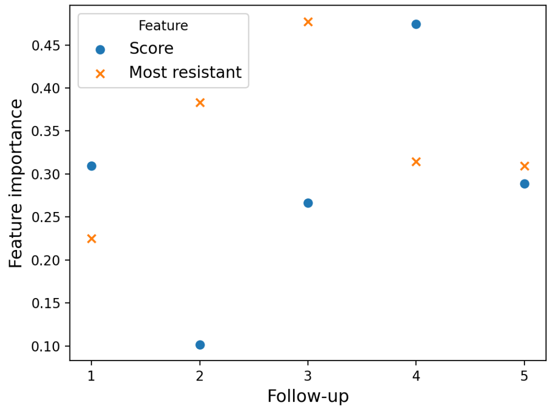


Figure S4: **Clinical response prediction features.** Importance assigned to the *score* or *resistant* features when training a Random Forest using the features from scClone2DR for predicting clinical response.

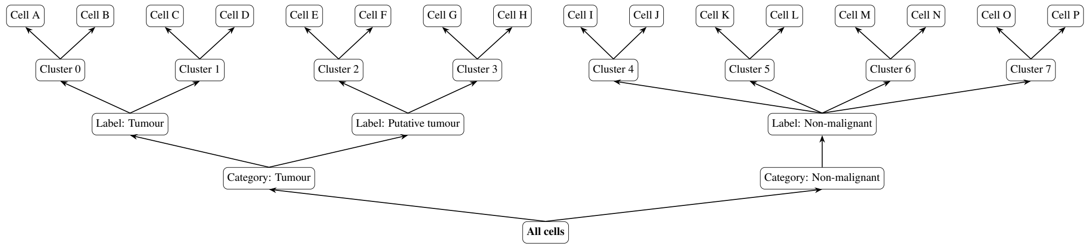


Figure S5: **AML cell hierarchy.** Illustrative example of the hierarchical assignments of metacells for the AML analysis.

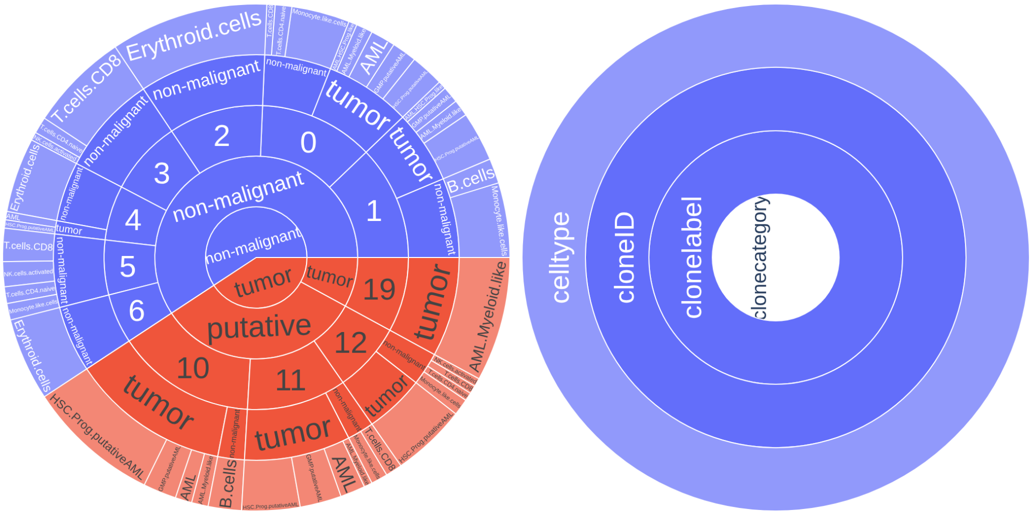


Figure S6: **AML clone assignment.** Example clonal assignment and labelling of an AML sample.

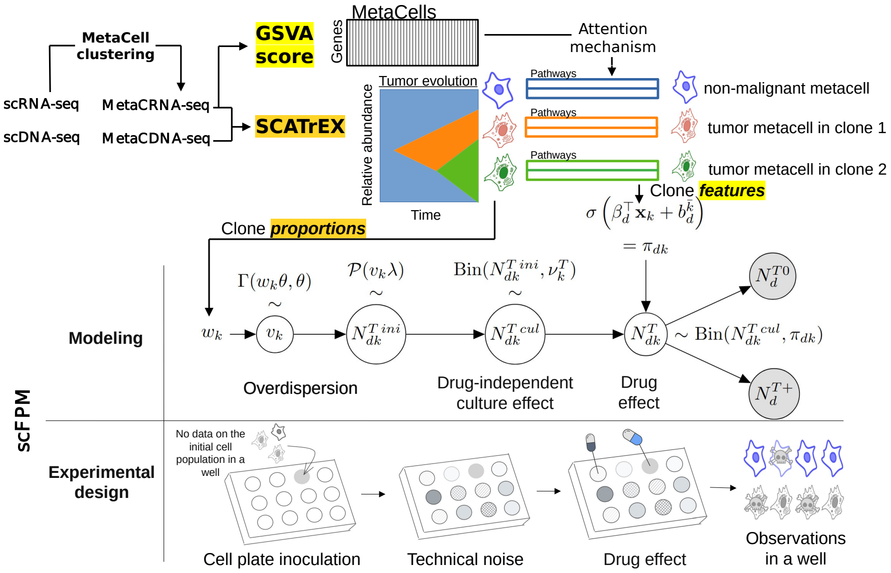


Figure S7: **Technical overview of scClone2DR.** The input scDNA-seq data provides an evolutionary history of the tumour and its clones onto which the scRNA-seq data is mapped. The scRNA-seq data is processed as metacells for which pathway scores are extracted (as depicted). Alternatively, for a reduced set of genes, RNA features are embedded into a lower-dimensional space through a VAE. The features pass through an attention mechanism to provide the clonal features for each tumour clone and for the non-malignant cells. The clonal features enter the scClone2DR model for drug effects, where the survival probability of the various clones is derived from the features and is inferred from the scFPM ex-vivo drug screening data where we only observe the final number of surviving non-malignant and tumour cells. The scClone2DR model also accounts for experimental noise in the drug screening, including overdispersion, and confounding factors like well position and cell density. The mathematical notation is explained in Supplementary Table S3 and here for clarity we have omitted the subscripts specifying the patient and replicate. The output is the predicted drug response of each tumour clone for each drug and for each patient.

### Supplementary Tables

Table S1: Overview of drugs with highest delta LOR per sample and clone across the samples of the Tumor Profiler melanoma cohort.

| sample | drug set | number of clones | number of drugs |
|---|---|---|---|
| MACEGEJ | Dabrafenib,Temozolomide | 5 | 2 |
| MACYHYS | Carboplatin,Dabrafenib,Erlotinib | 3 | 3 |
| MADEGOD | Dasatinib,Everolimus | 5 | 2 |
| MADIBUG | Carboplatin,Docetaxel | 6 | 2 |
| MAHACEB | Dabrafenib,Erlotinib,Olaparib,Paclitaxel,Temozolomide | 6 | 5 |
| MAHEFOG | Carboplatin,Dabrafenib,Dasatinib,Paclitaxel | 5 | 4 |
| MAJEFEV | Erlotinib,Everolimus,Paclitaxel | 6 | 3 |
| MAJOFIJ | Cisplatin,Erlotinib,Everolimus | 8 | 3 |
| MAKYGIW | Carboplatin,Temozolomide | 5 | 2 |
| MALYLEJ | Carboplatin,Dabrafenib,Erlotinib | 7 | 3 |
| MANOFYB | Alpelisib,Docetaxel | 3 | 2 |
| MAPOXUB | Docetaxel,Temozolomide | 3 | 2 |
| MECUGUH | Erlotinib,Everolimus,Temozolomide | 6 | 3 |
| MECYGYR | Alpelisib,Carboplatin,Dabrafenib,Temozolomide | 6 | 4 |
| MEDYCAR | Docetaxel,Erlotinib,Everolimus,Paclitaxel,Sorafenib,Temsirolimus | 6 | 6 |
| MEFOCUR | Dabrafenib,Everolimus,Gefitinib | 7 | 3 |
| MEHUFEF | Carboplatin,Cisplatin,Temozolomide,Temsirolimus | 7 | 4 |
| MEHYLOB | Carboplatin,Dabrafenib,Docetaxel,Temozolomide | 6 | 4 |
| MEKAKYD | Carboplatin,Dabrafenib,Temsirolimus | 4 | 3 |
| MEKOBAB | Docetaxel,Everolimus,Paclitaxel,Palbociclib | 4 | 4 |
| MELIPIT | Gefitinib,Sorafenib,Temozolomide | 6 | 3 |
| MELUKUZ | Carboplatin,Sorafenib,Temsirolimus | 8 | 3 |
| MELYPEB | Carboplatin,Dabrafenib,Olaparib,Paclitaxel,Temozolomide,Temsirolimus | 8 | 6 |
| MIFIMAB | Temozolomide | 5 | 1 |
| MIFOGIL | Everolimus,Temozolomide | 5 | 2 |
| MIGEKUT | Dabrafenib,Everolimus | 6 | 2 |
| MIGOFIW | Carboplatin,Paclitaxel,Temozolomide | 6 | 3 |
| MIJUCYK | Carboplatin,Temozolomide | 6 | 2 |
| MIKOBID | Carboplatin,Everolimus,Temsirolimus | 4 | 3 |
| MIKYCYN | Dasatinib,Erlotinib,Sorafenib,Temozolomide | 4 | 4 |
| MILABYL | Dabrafenib,Everolimus | 6 | 2 |
| MIMANAR | Alpelisib,Docetaxel,Gefitinib | 5 | 3 |
| MISYPUP | Carboplatin,Docetaxel,Erlotinib,Paclitaxel,Temozolomide | 6 | 5 |
| MOBUBOT | Docetaxel,Erlotinib,Gefitinib,Olaparib | 8 | 4 |
| MOCELOJ | Carboplatin,Dabrafenib,Sorafenib,Temozolomide | 9 | 4 |
| MODUDOL | Carboplatin,Dasatinib,Erlotinib | 8 | 3 |
| MOFYCAG | Dabrafenib,Erlotinib | 3 | 2 |
| MOGYLIP | Carboplatin,Olaparib | 3 | 2 |
| MOTAMUH | Carboplatin,Dabrafenib,Sorafenib | 7 | 3 |
| MOVAZYQ | Alpelisib,Carboplatin,Dabrafenib,Everolimus,Olaparib,Paclitaxel,Sorafenib | 9 | 7 |
| MUBOMEF | Everolimus,Paclitaxel | 6 | 2 |
| MUCADOP | Carboplatin,Paclitaxel | 5 | 2 |
| MUGAKOL | Dabrafenib,Olaparib,Sorafenib | 6 | 3 |
| MUKAGOX | Dabrafenib,Temozolomide | 6 | 2 |
| MYBYHER | Carboplatin,Erlotinib,Temozolomide | 5 | 3 |
| MYJILAS | Carboplatin,Olaparib,Paclitaxel | 5 | 3 |
| MYKOKIG | Carboplatin,Dasatinib,Olaparib | 4 | 3 |
| MYNELIC | Erlotinib,Gefitinib,Olaparib | 5 | 3 |

| Comparison | $p$-value |
|---|---|
| scClone2DR vs Clonal FM | 0.14 |
| scClone2DR vs Baseline | $2.2 \times 10^{-3}$ |
| scClone2DR vs scFPM | $1.5 \times 10^{-3}$ |
| Clonal FM vs Baseline | $3.3 \times 10^{-2}$ |
| Clonal FM vs scFPM | 0.041 |
| Baseline vs scFPM | 0.45 |

Table S2: Pairwise DeLong tests comparing ROC AUC values for the clinical response modelling shown in Figure 5.

Table S3: List of notation.

| | |
|---|---|
| **Indices:** | |
| $R_T$ | Number of wells with treatment ($r \in \{1, \dots, R_T\}$) |
| $R_C$ | Number of control wells ($r \in \{1, \dots, R_C\}$) |
| $I$ | Number of samples ($i \in \{1, \dots, I\}$) |
| $D$ | Number of drugs ($d \in \{1, \dots, D\}$) |
| $K_i$ | Number of tumour clones for sample $i$ |
| $L$ | Dimension of the feature vectors |
| **Parameters:** | |
| $\beta_d$ | Regression coefficients for drug $d$ |
| $b_d^0$ | Offset coefficient for drug $d$ for the clone with non-malignant cells |
| $b_d^+$ | Offset coefficient for drug $d$ for clones with tumour cells |
| $w_{ik}$ | Proportion of cells in clone $k \in \{0, \dots, K_i\}$ for sample $i$ |
| $\theta_i$ | Overdispersion parameter for sample $i$ for drug data |
| $\theta^{rna}$ | Overdispersion parameter for the single-cell RNA data |
| $\nu_{ir}^{C0}$ | Survival probability of non-malignant cells for patient $i$ in the $r$-th control well |
| $\nu_{ir}^{C+}$ | Survival probability of tumour cells for patient $i$ in the $r$-th control well |
| $\nu_{idr}^{T0}$ | Survival probability of non-malignant cells for patient $i$ in the $r$-th well treated with drug $d$ |
| $\nu_{idr}^{T+}$ | Survival probability of tumour cells for patient $i$ in the $r$-th well treated with drug $d$ |
| $\pi_{idk}$ | Survival probability of cells in clone $k$ for sample $i$ due to the effect of drug $d$ |
| $\sigma_0$ | Standard deviation for the prior distribution over $\gamma_0$ |
| $\sigma_+$ | Standard deviation for the prior distribution over $\gamma_+$ |
| $\mu, \Sigma$ | Mean and covariance matrices of the variational distribution on the random vector $(\gamma_0, \gamma_+)$ |
| $\eta$ | Coefficients of the GAM model used to obtain the culture survival probabilities |
| **Random variables:** | |
| $N_{ik}^{rna}$ | Number of cells in clone $k \in \{0, \dots, K_i\}$ for sample $i$ for RNA data |
| $N_{idr}^{T0}$ | Number of non-malignant cells for sample $i$ in well $r$ with drug $d$ |
| $N_{idr}^{T+}$ | Number of tumour cells for sample $i$ in well $r$ with drug $d$ |
| $N_{idr}^{T}$ | Total number of cells for sample $i$ in well $r$ with drug $d$ |
| $N_{ir}^{C0}$ | Number of non-malignant cells for sample $i$ in control well $r$ |
| $N_{ir}^{C+}$ | Number of tumour cells for sample $i$ in control well $r$ |
| $N_{ir}^{C}$ | Total number of cells for sample $i$ in control well $r$ |
| $\gamma_0$ | Vector used to obtain the attention weights which are then applied to derive the feature vectors $\mathbf{x}_{i0}$ for the groups of non-malignant cells. |
| $\gamma_+$ | Vector used to obtain the attention weights which are then applied to derive the feature vectors $\mathbf{x}_{ik}$ for any clone of tumour cells. |
| **Glossary:** | |
| clusters | Groups of cells obtained using only a single modality: scRNA-seq |
| clones | Groups of cells obtained using multiple modalities: scDNA-seq and scRNA-seq. |

### Supplementary Material

#### S1 Simulations

To generate simulated data, by default we set parameters to approximate the settings for the real data. For each patient, we consider 6 tumour clones (i.e. $K = K_i = 6$ for all $i$). The total number of control wells per patient is set to $R_C = 24$. For all patient and all drug, the number of replicates for treated wells is set to 5 (i.e. $R_T = 5$). The total number of patients is 100 with typically 50 patients for training and 50 patients in the test set.

The design matrix $\mathbf{X}$ is then created with dimensions $(K+1) \times N \times L$. Each entry is obtained as the absolute value of a sample from a Gaussian distribution with mean 0 and variance 0.3, scaled by $1 - k/K$ to decrease linearly across clones.

The design matrix $\mathbf{X}$ is then provided to the user. By doing so, we do not need to rely on the attention mechanism described above to get feature vectors at the clone level (since they are already given at this resolution). This way, we avoid the latent random variables $\gamma_0, \gamma_+$ and directly compute the maximum likelihood estimators without using variational inference. For any drug $d$, the entries of $\beta_d$ are obtained by taking the absolute values of i.i.d. samples from a Gaussian distribution with mean 0 and variance $1/L$. Overdispersion parameters $\theta_i$ are set by default to $10^3$. The design matrices for the assay effect – represented by $\mathbf{x}_{ir}^C$ and $\mathbf{x}_{idr}^T$ – are one dimensional and sampled from a Gaussian distribution centred around $\log(0.6/0.9)$ with variance 0.03 and $\eta = 1$ responses.

We additionally generated simulated data in an easier setting with more replicates for treated wells (20 compared to the 5 previously) and with less overdispersion ($\theta = 10^5$ compared to the previous $10^3$).

**Comparison methods** We compare the performance of scClone2DR to the following alternatives:

***Baseline:*** We define a simple baseline model where we consider only two groups of cells (one clone for the non-malignant cells and one for the tumour cells) and where the drug-specific survival probabilities for each clone are shared across patients and do not depend on any features of the clone. These parameters are learned from the data, and the baseline model is a specific instance of our scClone2DR model where we set $\beta_d = \mathbf{0}$ for all drugs $d$ in Equation (1).

***Bulk:*** We also define another simple model where we imagine we only had bulk measurements in that we compute a single feature vector for each patient by averaging over all clones: $\mathbf{x}_i := \sum_{k=0}^{K_i} w_{ik}\mathbf{x}_{ik}$. In this setting we would not have access to the clonal fractions as measured from the combined scDNA and scRNA analysis. Consequently, the graphical model from Supplementary Figure S17 is simplified to include only the two branches related to control and treated wells. In our modelling, we continue to consider two group of cells, namely the group of non-malignant cells and the group of tumour cells, for which we want to estimate the proportions $w_{i0}$ and $w_{i+} := \sum_{k \geq 1} w_{ik}$ as well as the survival probabilities

$$\pi_{id0} = \sigma\left(\mathbf{x}_i^\top \beta_d + b_d^0\right) \text{ and } \pi_{id+} = \sigma\left(\mathbf{x}_i^\top \beta_d + b_d^+\right).$$

***Dual bulk:*** As an additional variant we consider a pseudo-bulk approach where, based on cell-typing as performed with scRNA-seq data, we perform an average over non-malignant cells and over tumour cells separately. Thus, we consider the feature vectors $\mathbf{x}_{i0}$ and $\mathbf{x}_{i+} := \sum_{k \geq 1} \frac{w_{ik}}{1-w_{i0}} \mathbf{x}_{ik}$.

To define further alternatives we consider the main output of the ex-vivo drug screening which is the fraction of non-malignant cells in control wells $f_i^{C0} := \frac{N_i^{C0}}{N_i^C}$ (or in treated wells $f_{id}^{T0} := \frac{N_{id}^{T0}}{N_{id}^T}$). Using a first-order approximation of the expected values of these quantities

$$\mathbb{E}[f_i^{C0}] \approx \frac{\mathbb{E}[N_i^{C0}]}{\mathbb{E}[N_i^C]} = w_{i0}, \quad \text{and} \quad \mathbb{E}[f_{id}^{T0}] \approx \frac{\mathbb{E}[N_{idr}^{T0}]}{\mathbb{E}[N_{id}^T]} = \frac{\pi_{id0} w_{i0}}{\sum_{k=0}^{K_i} \pi_{idk} w_{ik}},$$

we obtain that

$$\frac{\mathbb{E}[f_i^{C0}]}{\mathbb{E}[f_{id}^{T0}]} \approx \sum_{k=0}^{K_i} w_{ik} \frac{\pi_{idk}}{\pi_{id0}}. \tag{2}$$

Based on Equation (2), we propose two further alternatives to model clonal drug response through differently parametrizing the ratios of survival probabilities $\frac{\pi_{idk}}{\pi_{id0}}$. In each scenario, we use the proxies $\bar{w}_{ik}$ for $w_{ik}$ where $\bar{w}_{ik}$ is derived from the proportion of cells within each clone calculated from the combined scDNA and scRNA analysis, $\left(w_{ik}^{RNA}\right)_k$. Adjustments are made for the proportion of non-malignant cells observed in control wells through averaging across all replicates.

We will denote by $f_{ir}^{C0} := \frac{N_{ir}^{C0}}{N_{ir}^{C}}$ (or $f_{idr}^{T0} := \frac{N_{idr}^{T0}}{N_{idr}^{T}}$) the fraction of non-malignant cells for patient $i$, in the $r$-th control-well replicate (or in the $r$-th well replicate corresponding to treatment with drug $d$). Model parameters are learned by minimizing, with respect to $\theta$, the quadratic loss

$$\sum_i \sum_d \left( y_{id} - \sum_{k=0}^{K_i} \bar{w}_{ik} f_\theta(\mathbf{x}_{ik}) \right)^2,$$

where $y_{id} := \frac{\frac{1}{\#}\sum_r f_{ir}^{C0}}{\frac{1}{\#}\sum_r f_{idr}^{T0}}$ is the ratio of the observed fractions of non-malignant cells in control and treated wells, and where $f_\theta$ is the model with parameter $\theta$ used to output the predicted ratio of survival probabilities $\frac{\pi_{idk}}{\pi_{id0}}$.

***Neural Network:*** We consider an off-the-shelf machine learning neural network approach. Namely, we implement a feed-forward neural network with one hidden layer of dimension $128 \times 64$ with ReLU activation functions and we apply the exponential map at the end of the output layer to ensure the required positivity. The input of our model is the vector obtained by concatenating the feature vector of the patient and the true feature vector of the drug, namely $(\mathbf{x}_{ik}^\top, \beta_d^\top)$.

***Factorization Machine:*** Finally, we include a factorization machine model for drug response prediction that can be understood as the counterpart of the scDrug tool [32] repurposed for our framework. The ratio of survival probabilities are modelled as follows

$$\forall k \geq 1, \quad \log\left(\frac{\pi_{idk}}{\pi_{id0}}\right) = c_d + \mathbf{q}_d^\top \mathbf{W} \mathbf{x}_{ik},$$

where $c_d$ is an offset parameter specific to drug $d$, $\mathbf{q}_d$ is a latent representation of drug $d$, while the matrix $\mathbf{W}$ is a transformation that projects clone feature $\mathbf{x}_{ik}$ onto the latent space.

For both the Factorization Machine and Neural Network models, clone proportions were initially derived from the RNA data (as combined with the DNA analysis) and subsequently adjusted by reweighting the non-malignant and tumour clones to ensure that the overall tumour cell fraction aligns with observations from control wells. This correction addresses potential biases introduced by drug-independent culture effects.

**Performance metrics** We consider the following performance metrics to highlight different aspects of the models performances.

***KL-divergences:*** First, we calculate the average KL-divergence across all test patients between the estimated clone proportions $\hat{w}_{ik}$ and the true proportions $w_{ik}$ for both our approach and the baseline model. This metric is not considered for the other methods, as they either estimated proportions from the scDNA and scRNA data (FM and NN models), or consider only two groups of cells (bulk and dual bulk models). Next, we assess the accuracy of the estimated survival probabilities by computing the average KL-divergence across test patients, drugs, and clones between the estimated survival probabilities $\hat{\pi}_{idk}$ and the true probabilities $\pi_{idk}$.

***$L^1$-errors:*** Along with the global accuracy between simulated and inferred parameters measured by the KL-divergence, we focus more on the largest errors for more clinically-relevant quantities. First, across all patients and drugs, we evaluate the $L^1$ error between the estimated overall survival rate $\sum_{k\geq 0} \hat{w}_{ik}\hat{\pi}_{idk}$ and the true one $\sum_{k\geq 0} w_{ik}\pi_{idk}$. Another relevant quantity is the drug effect $e_{id}$ defined as the ratio of the fraction of tumour cells with drug $d$ and the one without treatment, namely

$$e_{id} := \frac{1}{1-w_{i0}} \frac{\sum_{k\geq 1} w_{ik}\pi_{idk}}{\sum_{k\geq 0} w_{ik}\pi_{idk}}. \tag{3}$$

We compute the $L^1$ error between the estimated drug effects and the true ones.

***Drug scores:*** We further consider drug scores measuring how effective a drug is at treating the entire tumour. As such we define a score $s_{id}$ for each patient-drug pair $(i, d)$ as the ratio of the survival probability of non-malignant cells to that of the most resistant tumour clone:

$$s_{id} := \frac{\pi_{id0}}{\max_{k\geq 1} \pi_{idk}}. \tag{4}$$

Thus, the larger $s_{id}$, the more effective the drug. As a summary, we compute the average Spearman correlation across all patients between the estimated scores $(\hat{s}_{id})_d$ and the true scores $(s_{id})_d$. For those models that handle all tumour clones, we also evaluate their ability to rank tumour cell groups from most to least resistant to the drug. We compute the average Spearman correlation across all patients and drugs between $\left(\frac{\hat{\pi}_{id0}}{\hat{\pi}_{idk}}\right)_{k\geq 1}$ and $\left(\frac{\pi_{id0}}{\pi_{idk}}\right)_{k\geq 1}$.

***Fold-changes:*** The fold change is defined as the difference between the average (over all replicates) of the log–fraction of non-malignant cells in control wells and that in wells treated with drug

$$\frac{1}{R_C}\sum_{r=1}^{R_C} \log(\mathbb{E}[f_{ir}^{C0}]) - \frac{1}{R_T}\sum_{r=1}^{R_T} \log(\mathbb{E}[f_{idr}^{T0}]).$$

We compute this fold change using both the ground-truth parameters and those estimated by the different methods.

In Figure 2 we display the fold-changes (a) and drug scores (b), while we further report on summaries of the additional metrics in Supplementary Table S4. Results in the easier setting with more replicates and less overdispersion are presented in Supplementary Figure S1.

| | scClone2DR | Baseline | Dual bulk | Bulk | FM | NN |
|---|---|---|---|---|---|---|
| $\mathrm{KL}((\hat{w}_{ik})_k, (w_{ik})_k)$ | 0.111 | **0.100** | 0.103 | ✗ | ✗ | ✗ |
| $\mathrm{KL}((\hat{\pi}_{idk})_k, (\pi_{idk})_k)$ | **0.008** | 0.041 | 0.039 | 0.041 | 0.049 | 0.049 |
| $L^1$ error overall survival rate | **0.471** | 0.938 | 0.583 | 0.916 | 2.363 | 2.163 |
| $L^1$ error on drug effects | 0.011 | 0.022 | **0.010** | 0.022 | 0.053 | 0.056 |
| Spearman on drug scores | **0.699** | 0.083 | 0.593 | 0.297 | 0.038 | -0.369 |
| Spearman on clone scores | 0.936 | ✗ | ✗ | ✗ | **0.947** | 0.657 |

Table S4: Comparison of scClone2DR with benchmark methods over various evaluation metrics.

### S2 Further scClone2DR simulation results

**Consistency of the estimators** We assess the consistency of the Maximum Likelihood Estimation (MLE) estimates by increasing the number of replicates for treated wells and by considering different level of overdispersion. Supplementary Figure S8a illustrates the Kullback-Leibler divergence between the estimated proportions and their true counterparts. Additionally, we depict the evolution of the Mean Squared Error (MSE) between the true regression coefficients and the estimated values (Supplementary Figure S8b). These findings demonstrate our ability to accurately recover the true proportions and regression coefficients, including the genuine survival probabilities, when a sufficient number of replicates is available (Supplementary Figure S9). These plots also provide a means to quantify the impact of overdispersion: the larger $\theta$, the smaller the overdispersion, and consequently, the smaller the estimation error on the parameters of interest. As well as examining the errors, we also visualise the learned parameters compared to the true ones. Supplementary Figure S10a shows the learned proportions and the ones used to generate the data for some specific patients in the test set. In Supplementary Figure S10b, the learned regression coefficients $\beta_d$ are compared to the true coefficients for some drugs.

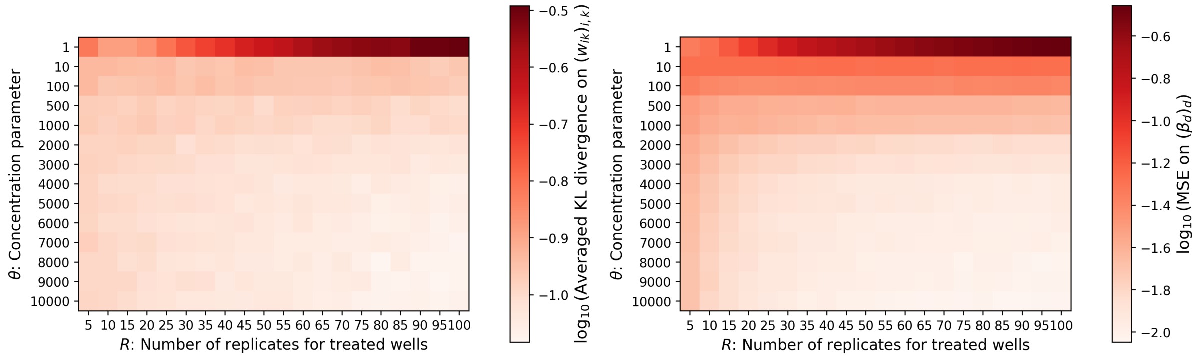


(a) Consistency of the estimated clone proportions. (b) Consistency of the estimated regression coefficients.

Figure S8: (a) KL divergence between the true clone proportions and the estimated ones. (b) Mean squared error between the true regression coefficients $(\beta_d)_{d\in[D]}$ and the estimated ones. For each plot, we average the errors across all patients and over 40 estimates obtained from different random generated datasets.

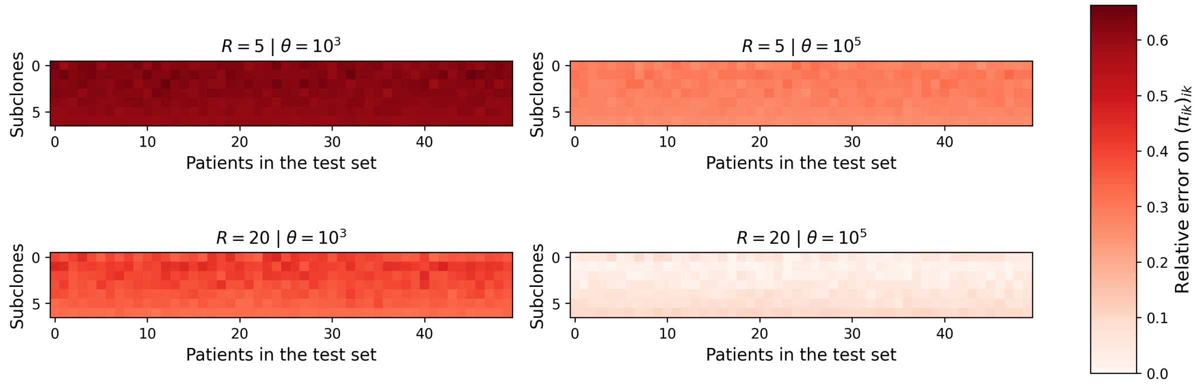


Figure S9: Relative errors on the survival probability for the drug n°1 on the test set.

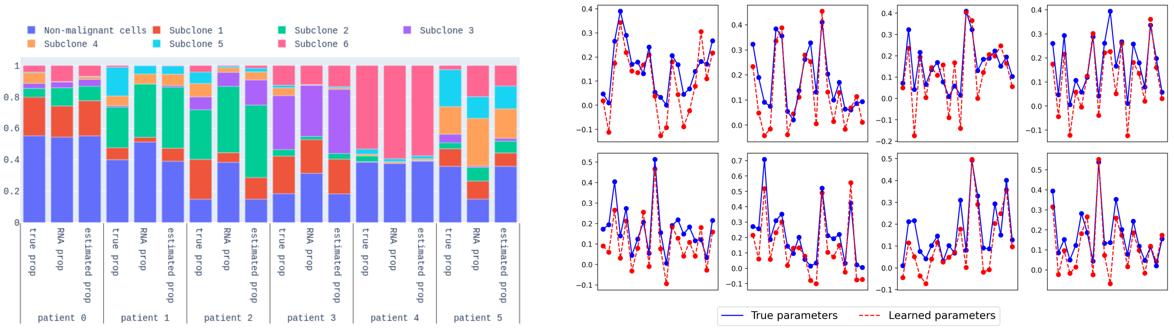


(a) Consistency of the estimated clones proportions. (b) Consistency of the estimated regression coefficients.

Figure S10: (a) For six patients from the test set, we show the true proportions of each clone, the ones observed from the RNA data and the ones estimated by our model. (b) For eight drugs, we show the true regression coefficients $\beta_d$ and the estimated ones. The plots are obtained considering a dataset generated with $R = 100$ replicates for treated wells and a concentration parameter $\theta = 10^4$.

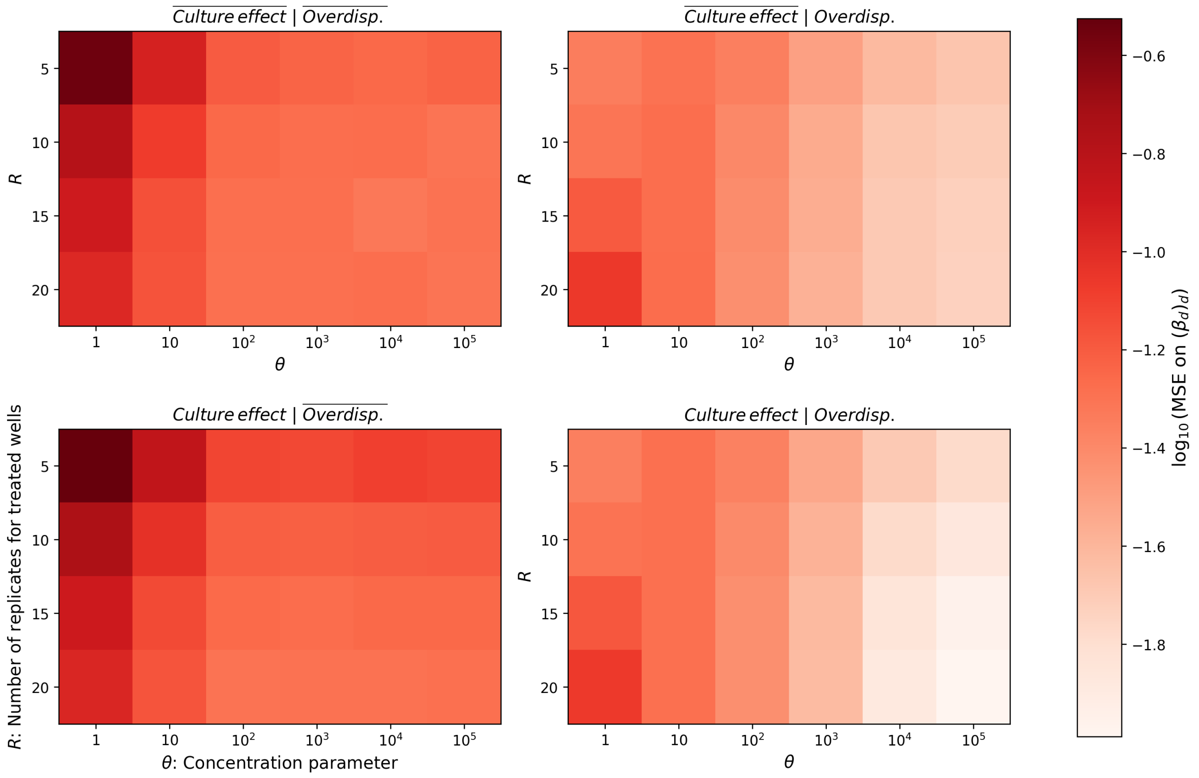


Figure S11: MSE on the different drug regression coefficients (averaging across all samples). We use four different methods: learning a culture effect ($Culture\,effect$) or not ($\overline{Culture\,effect}$), and modeling overdispersion ($Overdisp.$), or considering only multinomial distributions ($\overline{Overdisp.}$).

**Ablation study** In scClone2DR we model the overdispersion exhibited by the real data. This overdispersion can be primarily attributed to the way cells are put in suspension before being profiled with the different technologies. Consequently, we account for the overdispersion effect not only in the drug data but also in the scRNA-seq data (Supplementary Section S6). We also model effects from the experimental design which is mimicked in the simulated data.

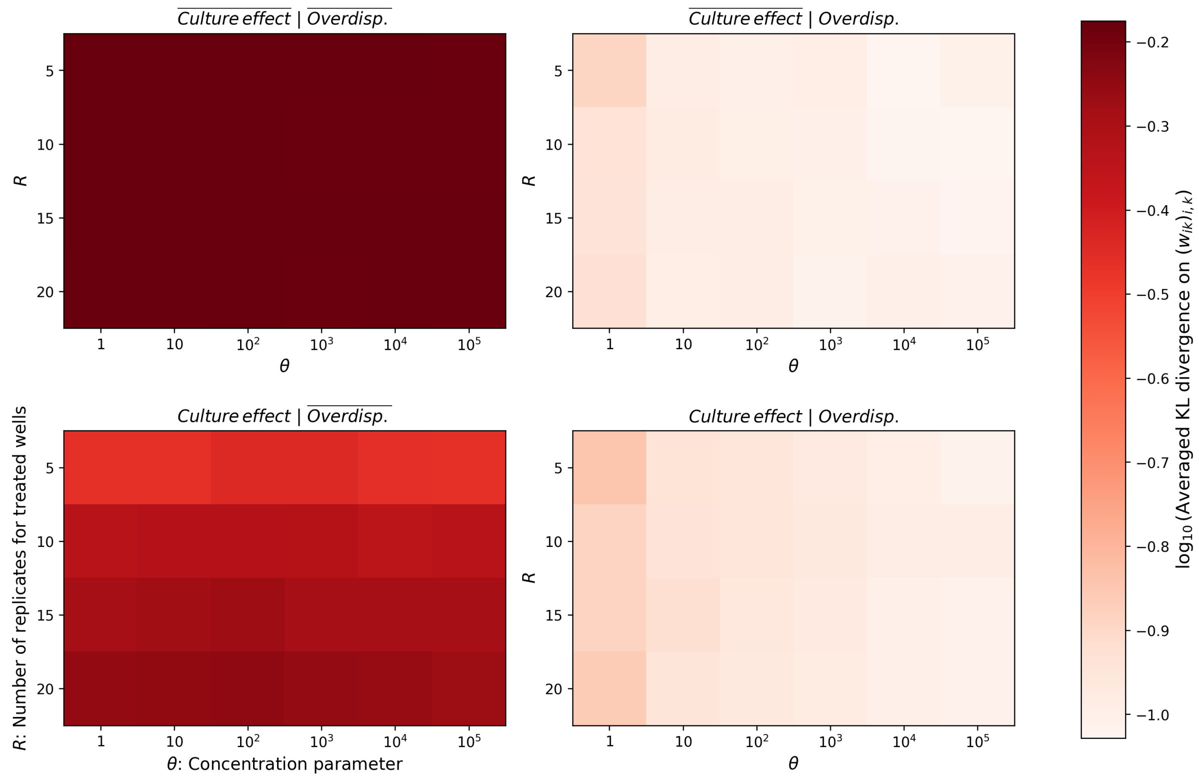


Figure S12: KL divergence on the clone proportions (averaging across all samples). We use four different methods: learning a culture effect ($Culture\,effect$) or not ($\overline{Culture\,effect}$), and modelling overdispersion ($Overdisp.$), or considering only multinomial distributions ($\overline{Overdisp.}$).

To quantify the importance of modelling both the overdispersion and experimental effects we conduct ablation studies. We generate simulated data by employing the generative model described in Supplementary Section S6 and selecting the overdispersion parameters in a manner that closely mimics the behaviour of the real data. We then can choose to 1) model a correction term for the experimental biases or not, and/or 2) to include/exclude the overdispersion effects from our model. While 1) consists in fixing all parameters modelling the culture effect ($\nu_{idr}^{T0}, \nu_{idr}^{T0}, \nu_{ir}^{C0}, \nu_{ir}^{C+}$) to the same constant value, 2) means that the $\theta_i$ go to $+\infty$ in our model (equivalently we replace the Dirichlet-multinomial/beta-binomial distributions with multinomial/binomial distributions respectively).

We conducted this ablation study for different simulated dataset, considering different overdispersion level and different number of replicates for treated wells. We observe a strong drop in performance when not modelling the experimental and/or overdispersion effects both in terms of the MSE (Supplementary Figure S11) and the KL-divergence (Supplementary Figure S12), with the overdispersion modelling having the stronger effect.

**Reliability** Though in simulations we can observe the accuracy and performance of scClone2DR in recovering the underlying parameters, as above, one of the challenges in assessing its reliability for use on real data is that we only observe some quantities and not all parameters. To explore this, we focus on the fold change which quantifies the effect of a drug on cells from a specific patient. The fold change can be either estimated using the observed data, or derived from our generative model with either the learned parameters or the true ones from the simulation setting. In all cases, the fold change is defined as the average (over all replicates) of the log of the fraction of non-malignant cells in control wells minus the average (over all replicates) of the log of the fraction of non-malignant cells in treated wells. Namely for patient $i$ and drug $d$, we have:

$$\text{Observed fold change} = \frac{1}{\#}\sum_r \log(f_{ir}^{C0}) - \frac{1}{\#}\sum_r \log(f_{idr}^{T0}),$$

$$\text{Predicted fold change} = \frac{1}{\#}\sum_r \log(\widehat{f_{ir}^{C0}}) - \frac{1}{\#}\sum_r \log(\widehat{f_{idr}^{T0}}),$$

$$\text{True fold change} = \frac{1}{\#}\sum_r \log(\overset{*}{f}{}_{ir}^{C0}) - \frac{1}{\#}\sum_r \log(\overset{*}{f}{}_{idr}^{T0}),$$

where $\widehat{f_{ir}^{C0}}$ (or $\overset{*}{f}{}_{ir}^{C0}$) is the expected value of the fraction of non-malignant cells for patient $i$ in the $r$-th control well and $\widehat{f_{idr}^{T0}}$ (or $\overset{*}{f}{}_{idr}^{T0}$) is the expected value of the fraction of non-malignant cells for patient $i$ in the $r$-th well with treatment drug $d$ considering our generative model with the learned parameters (or the true parameters).

We consider two settings: an easier setting with a large number of replicates for treated wells and a small overdispersion effect, and another more realistic (and more challenging) setting where both overdispersion and the number of replicates (R=5) mimic those found in real data.

When we compare the predicted fold change to the true simulated values, scClone2DR successfully recaptures the underlying effects (Supplementary Figure S13, right column), as in Figure 2a for the more challenging setting. However, when we compare to the observed value we only see good agreement for the easier setting (Supplementary Figure S13f). In the more realistic setting, the noise in the real data is such that we see little correlation between the predicted and observed values (Supplementary Figure S13c). As scClone2DR effectively accounts for this noise during the learning phase to accurately estimate parameters, we are able to recover the ground truth (Supplementary Figure S13a) even when these are not directly evident in the observed data. In general though, we must exercise caution in using observed data alone to evaluate model performance since this could be misleading in high noise settings.

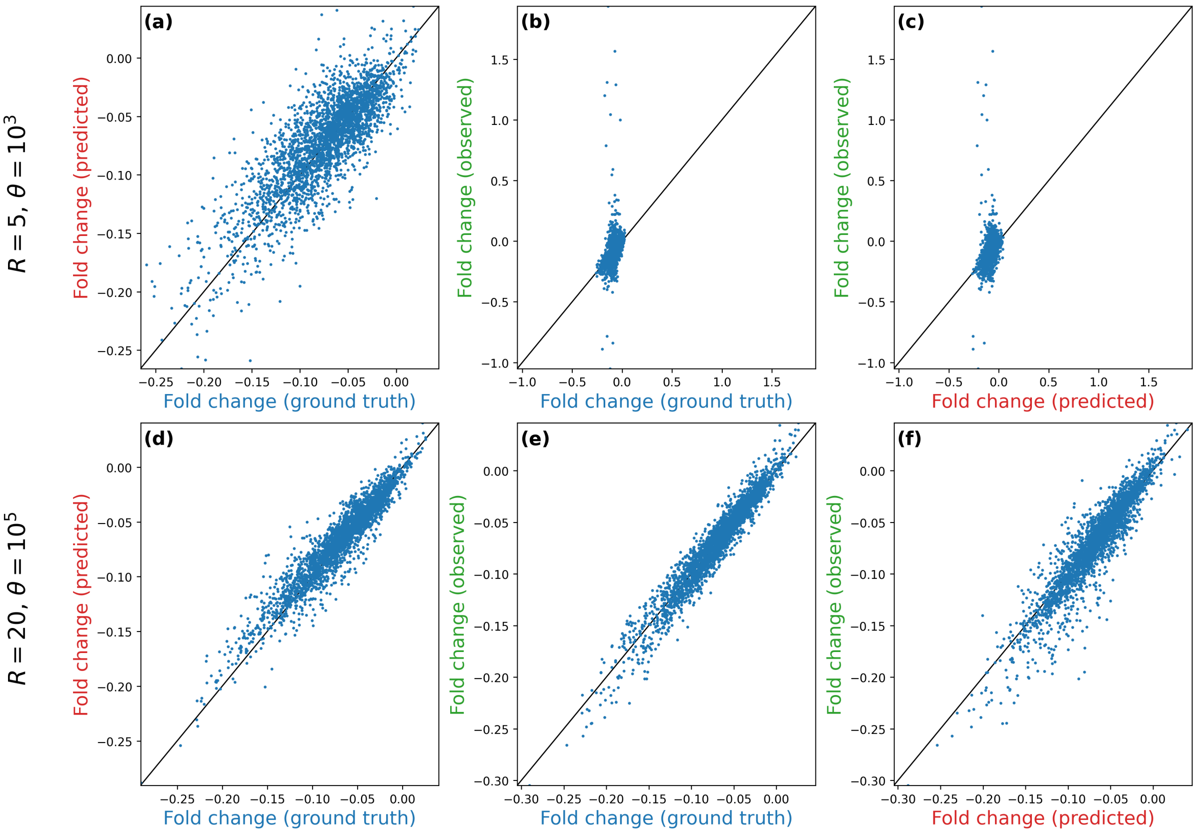


Figure S13: We consider two different dataset, one generated with $R = 5$ and $\theta = 10^3$ (top row) and one generated with $R = 20$ and $\theta = 10^5$ (bottom row). We show the predicted fold change versus the one computed using the true parameters (left column), the observed fold change versus the one computed using the true parameters (middle column), and the observed fold change versus the predicted one (right column).

### S3 Additional analyses on the melanoma cohort

**scClone2DR inference is stable** For the melanoma cohort we additionally assessed the stability of the inference. Since this is performed through a stochastic approach (Methods) there may be variations in results across different runs. Testing over 100 different random seeds, we observe a coefficient of variation (standard deviation divided by mean) of under 0.2 in 95% of cases. Those with larger coefficients of variation are concentrated on the less relevant regression coefficients, i.e. those with small means (Supplementary Figure S14a). With the sigmoid transformation in the generalised linear model (Methods) for the more relevant survival probability predictions the coefficient of variation is then under 5% in nearly all cases (Supplementary Figure S14b). Overall we observe high stability in our scClone2DR results.

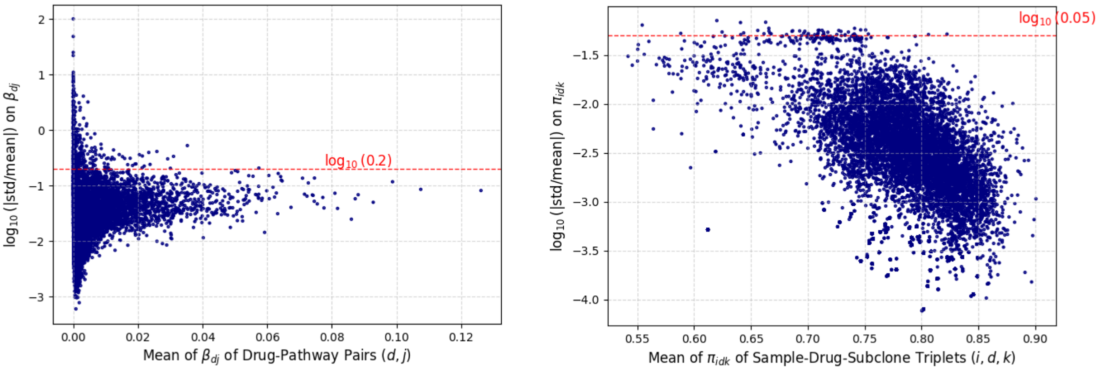


(a) (b)

Figure S14: Log of the ratio between the standard deviation and the mean (over 100 different random seeds) of the $\beta_{dj}$ regression coefficients (a) and the survival probabilities $\pi_{idk}$ (b) as a function of their mean values. The horizontal lines represent a coefficient of variation of 20% and 5% respectively.

**Uncertainty quantification with scClone2DR is well calibrated** We estimated the model parameters using a training set comprising 60% of the patients. By fixing the model parameters to their learned values, the generative nature of scClone2DR allows us to produce a complete predictive distribution of the tumour cell fractions for treated wells of held-out test patients. Specifically, we employed a model-based parametric bootstrap to simulate 100 realizations of tumour cell fractions for each drug-treated well. For a target coverage of $1 - \alpha$, we generated a prediction interval for each (patient, well) pair by calculating the $\alpha/2$ and $1 - \alpha/2$ quantiles of the predictive distribution. To assess model calibration, we computed the empirical coverage, defined as the proportion of test pairs (patient, well) for which the observed tumour cell fraction fell within the predicted interval. For a perfectly calibrated model, this proportion should equal $1 - \alpha$. We find strong agreement (Supplementary Figure S15), with only a slight underestimation at very high coverages. This indicates that scClone2DR effectively models both drug effects and noise well and is appropriately calibrated.

### S4 Melanoma clinical response modelling

Longitudinal clinical data from the Melanoma cohort of the Tumor Profiler Study include multiple follow-up visits per patient. The number of follow-ups varies across individuals, typically ranging from two to six, with visits scheduled at approximately three-month intervals. Key clinical outcomes included treatment response, categorized into six groups: relapse-free, complete response, partial response, stable disease, progressive disease, and death. The response data is visualised in Supplementary Figure S16.

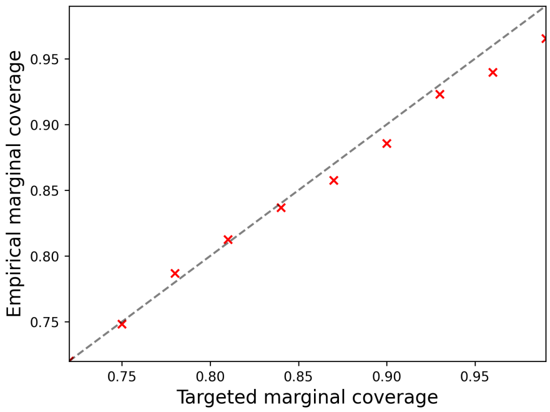


Figure S15: Testing the calibration of scClone2DR we compare the targeted coverage of prediction intervals on held-out data with the empirical coverage (red crosses). 60% of the data was used for training and results are averaged across all drugs (red crosses). Perfect calibration would correspond to the $y = x$ line (black dashed line).

**Survival modelling** Treatment response categories are highly imbalanced and unevenly distributed across follow-up visits, which limits the feasibility and stability of more complex longitudinal or ordinal models [49]. Moreover, such frameworks impose strong structural assumptions on the temporal evolution of clinical outcomes, which may not hold in the presence of multifactorial and treatment-dependent tumour dynamics. For these reasons, we adopted a simpler and more robust modelling strategy:

- ***Binary outcome formulation.*** We dichotomised the original six treatment response categories into two groups, combining relapse-free, complete response, and partial response into a single favourable-outcome class, and the remaining categories into an unfavourable-outcome class. This reduces class imbalance and centres the task on the most clinically meaningful distinction.
- ***Random Forest classifiers.*** We employed Random Survival Forests (akin to [50]) as they provide flexible, non-parametric estimators that require minimal assumptions about the underlying data-generating mechanism, making them well suited for heterogeneous cohorts and modest sample sizes.
- ***Independent models per follow-up.*** We trained a separate classifier at each follow-up time point. This avoids imposing rigid temporal dependencies, such as Markovian assumptions, on the disease trajectory and allows the relationship between features and outcomes to evolve freely across visits, reflecting the potentially shifting biological impact of different therapeutic interventions.

**Comparison methods** For each follow-up time point $t$, we trained separate Random Forest classifiers. The classifiers were trained using 500 decision trees with a maximum depth of 4 and a minimum of 3 samples per leaf. At each split, the number of candidate features was set to the square root of the total number of features. To account for class imbalance, class weights were set using a balanced weighting scheme, assigning weights inversely proportional to class frequencies in the training data. This procedure was applied across four different feature configurations.

- ***Baseline:*** This configuration includes patient age, sex, brain metastasis status, tumour stage, tumour mutational burden, immune infiltration status, presence of BRAFV600E mutation and subtype, as well as the number of days of immunotherapy or chemotherapy since the biopsy.

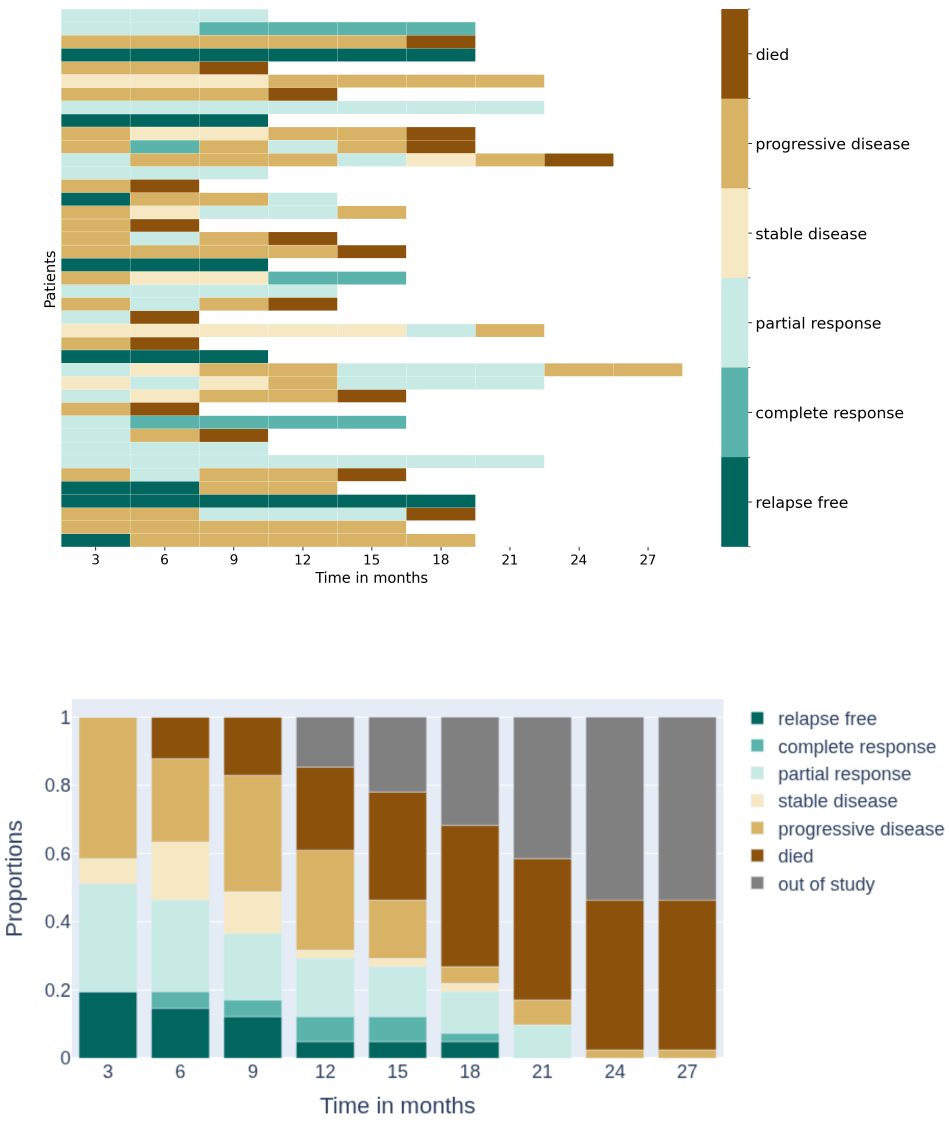


Figure S16: Visualization of the clinical response data for the Melanoma cohort.

- **FPM:** This model augmented these features with a score defined as

$$\mathrm{score}(t, w_{ik}^{\mathrm{FPM}}, \pi_{idk}^{\mathrm{FPM}}),$$

where relative survival probabilities between clones are used. Here, the survival probability for non-malignant cells $\pi_{id0}^{\mathrm{FPM}}$ is fixed at 1, while a single group of tumour cells is considered with $\pi_{id1}^{\mathrm{FPM}}$ computed as

$$\pi_{id1}^{\mathrm{FPM}} = \frac{f_{id}^{T+}}{f_{id}^{T0}} \times \frac{f_i^{C0}}{f_i^{C+}},$$

with $f_{id}^{T+}$ and $f_{id}^{T0}$ representing observed fractions of tumour and non-malignant cells (averaged over all replicates) in drug-treated wells, respectively, and $f_i^{C+}$, $f_i^{C0}$ their counterparts in control wells. The weights $w_{ik}^{\mathrm{FPM}}$ are defined accordingly, namely

$$w_{i0}^{\mathrm{FPM}} = \frac{f_i^{C0}}{f_i^{C+} + f_i^{C0}} \quad \text{and} \quad w_{i1}^{\mathrm{FPM}} = 1 - w_{i0}^{\mathrm{FPM}}.$$

- ***clonal FM:*** This configuration used baseline features together with score and resistant features calculated based on clone proportions $w_{ik}^{\mathrm{FM}}$, derived from the integrated scDNA and scRNA data analysis, and adjusted to match the proportions of tumour cells observed in control wells.
- ***scClone2DR:*** Lastly, the configuration combines baseline features with analogous clone score and resistant features using the general weights $w_{ik}$ and survival parameters $\pi_{idk}$ learned by scClone2DR.

**Feature definition** As features for predicting response, we consider the predicted aggregate response of the tumour, as well as that of the most resistant clone. With $\mathcal{D}_t$ denoting the set of drugs administered to patient $i$ between follow-ups $t-1$ and $t$, we have the following features:

- ***Aggregate score*** feature at time $t$:

$$\mathrm{score}(t, w_{ik}, \pi_{idk}) = \prod_{s=1}^{t} \frac{\sum_{k\geq 1} w_{ik} \prod_{d\in\mathcal{D}_s} \pi_{idk}}{w_{i0} \prod_{d\in\mathcal{D}_s} \pi_{id0}} = \frac{\#\text{tumour}(t)}{\#\text{non-malignant}(t)}.$$

- ***Resistant*** feature at time $t$:

$$\mathrm{resistant}(t, w_{ik}, \pi_{idk}) = \prod_{s=1}^{t} \frac{w_{i0} \prod_{d\in\mathcal{D}_s} \pi_{id0}}{w_{ik^*} \prod_{d\in\mathcal{D}_s} \pi_{idk^*}} = \frac{\#\text{non-malignant}(t)}{\#\text{most resistant clone}(t)},$$

  where

$$k^* = \arg\max_{k\geq 1} \prod_{s=1}^{t} \prod_{d\in\mathcal{D}_s} \pi_{idk}.$$

**Training and evaluation** Models are trained and tested in a leave-one-subject-out manner, where each patient is held out for testing while the others are used for training. We compute the receiver operating characteristic (ROC) curves derived from averaged predicted probabilities for each follow-up and each held-out patient. We additionally compute feature importance scores for the *score* and *resistant* features across follow-ups.

### S5 Attention mechanism to extract clone features

Let us denote by $\mathbf{Z}$ the 4 dimensional tensor of dimension $I \times (K+1) \times N_{mc} \times L$ where $I$ is the total number of patients, $K := \max_{i\in I} K_i$ is the maximum number of clones across patients (with $K_i$ the number per patient $i$), $N_{mc}$ is the maximum number of metacells in the different clones across patients, and $L$ is the dimension of the representations of the metacells. $\mathbf{Z}_{i,k,c,:}$ is the representation of the $c$-th cell in clone $k$ for patient $i$ (latent representations learned from scVI or the GSVA scores). In case clone $k$ of patient $i$ contains strictly less than $c$ metacells, all entries of $\mathbf{Z}_{i,k,c,:}$ are set to NaN. We standardize the feature matrix $\mathbf{Z}$ by centering and scaling each feature $\ell \in [L]$ using the mean and standard deviation computed across all metacells and patients.

For a given patient $i \in I$, the embeddings for the set of non-malignant cells and the one for any tumour clone $k \geq 1$ are respectively

$$[\mathbf{x}_{i0} \mid \gamma_0] = \mathrm{MSoftmax}\left(\mathbf{Z}_{i,0,:,:}\gamma_0\right)^\top \mathbf{Z}_{i,0,:,:} \quad \text{and} \quad [\mathbf{x}_{ik} \mid \gamma_+] = \mathrm{MSoftmax}\left(\mathbf{Z}_{i,k,:,:}\gamma_+\right)^\top \mathbf{Z}_{i,k,:,:},$$

where $\mathrm{MSoftmax}$ is the masked softmax operator defined by first setting to $-\infty$ any entry of the input vector that is NaN and by computing the usual softmax function on this modified input. By convention, we consider that $0 \times \text{NaN} = 0$. The parameters $\gamma_0$ and $\gamma_+$ are used to weight features of non-malignant and tumour cells, respectively, producing features for the group of non-malignant cells and tumour clones that are subsequently used to predict drug response. By considering separate parameters, the model can produce group-level representations that capture distinct signals relevant for predicting drug response. We consider the Bayesian priors $\gamma_0 \sim \mathcal{N}\left(\mathbf{0}, \sigma_0^2 \text{Id}\right)$ and $\gamma_+ \sim \mathcal{N}\left(\mathbf{0}, \sigma_+^2 \text{Id}\right)$, with $\gamma_0$ and $\gamma_+$ thus being latent random variables. The posterior distribution on this parameter is intractable and we use variational inference to approximate the posterior distribution. The variational distribution considered is a multivariate Gaussian on the joint distribution of the latent variables, namely

$$(\gamma_0, \gamma_+) \sim \mathcal{N}(\mu, \Sigma), \quad \mu \in \mathbb{R}^{2L}, \quad \Sigma \in \mathbb{R}^{2L \times 2L}. \tag{5}$$

To reduce the number of parameters required to define the variational distribution, we impose a low-rank plus diagonal structure on the covariance matrix $\Sigma$, with the rank set to 10.

Variational inference makes use of an upper-bound on the log likelihood, called the evidence lower bound (ELBO), defined by:

$$\text{ELBO}(y; \Phi, \mu, \Sigma) = \mathbb{E}_{q_{\mu,\Sigma}(\gamma_0,\gamma_+)}\left[\log\left(p_\Phi(\gamma_0, \gamma_+, y)\right) - \log\left(q_{\mu,\Sigma}(\gamma_0, \gamma_+)\right)\right],$$

where $q_{\mu,\Sigma}(\gamma_0, \gamma_+)$ is the variational distribution given by Equation (5). The variational parameters $\mu, \Sigma$ and model parameters $\Phi = \{(\beta_d, b_d^0, b_d^+)_d, (w_{ik})_{i,k}, (\theta_i)_i, \theta^{rna}, \eta, \sigma_0, \sigma_+\}$ are then learned by minimizing the upper bound on the penalized negative log-likelihood obtained using the ELBO:

$$\widehat{\Phi} \in \arg\min_{\Phi,\mu,\Sigma} \left\{ \frac{-1}{n_{obs}} \text{ELBO}(y; \Phi, \mu, \Sigma) + \sum_{d=1}^{D} \left(\lambda_1 \|\beta_d\|_1 + \lambda_2 \|\beta_d\|_2^2\right) + \lambda_3 \eta^\top S \eta \right\}, \tag{6}$$

where $n_{obs}$ is the total number of observations. We apply Elastic Net regularization to the drug-specific survival probability coefficients, $\beta_d$. This approach combines an $\ell_1$ penalty to promote sparsity with an $\ell_2$ penalty to improve stability in the presence of correlated features. The last penalty term $\eta^\top S \eta$ accounts for experimental effects and is described in Supplementary Section S7. All hyperparameters $\lambda_1$, $\lambda_2$ and $\lambda_3$ were selected via leave-one-out cross-validation, using the tumour fraction as the prediction target and choosing the values that minimize the cross-validated prediction error.

Optimizing the penalized ELBO for the entire dataset can be slow in practice and we use Stochastic Variational Inference (SVI) [51] to bypass this limitation. The variational distribution being a multivariate Gaussian, one can rely on the so-called reparametrization trick [52] to efficiently use stochastic gradient descent to optimize the variational parameters $\mu$ and $\Sigma$. Fitting the model on the full cohort of melanoma patients required approximately one minute using 8 CPU cores.

Hyperparameters are selected through cross validation of the correlation coefficient between the predicted and the true fraction of tumour cells.

### S6 Noise and graphical modelling

We will denote by $N_{ik}^{rna}$ the number of cells in clone $k \in [K_i]$ for the sample $i$ (where $K_i$ refers to the total number of tumour clones for sample $i$). The set of all non-malignant cells are placed in the same group of cells indexed by 0 and its size for sample $i$ is $N_{i0}^{rna}$. For convenience, we will refer to this group of non-malignant cells as clone 0. $w_{i0}$ is the fraction of non-malignant cells for sample $i$ while the tumour cells are divided into clones where $w_{ik}$ is the fraction of the total cells belonging to clone $k$. $\mathbf{x}_{ik}$ represents the feature vector for cells assigned to clone $k$ within sample $i$, obtained as described above by learning attention weights used to aggregate features at the cell level, following the method described in [36].

The underlying clonal fractions are not observed directly, but measured through the sequencing which is a noisy process which we model with overdispersion. Namely we set an overdispersion parameter $\theta^{rna}$ shared across all patients and scClone2DR models the observed data through a Dirichlet-multinomial distribution:

$$(N_{ik}^{rna})_{k\in\{0,\dots,K_i\}} \mid \sum_{k=0}^{K_i} N_{ik}^{rna} \sim \text{DirMulti}\left(\sum_{k=0}^{K_i} N_{ik}^{rna}\,,\ (\theta^{rna} w_{ik})_{k\in\{0,\dots,K_i\}}\right). \tag{7}$$

The larger $\theta^{rna}$, the smaller the overdispersion (and we approach a multinomial distribution).

The same cell suspension, with the same latent clonal fractions, are placed in the wells for the drug screening. We model the initial number of cells again with overdispersion through a Gamma-Poisson compound distribution which leads to a Beta-binomial. For each sample $i$, for each well $j$ we include an overdispersion parameter $\theta_i$ specific to patient $i$ and model the latent proportions of each clone $k$ as $v_{ijk} \sim \Gamma(w_{ik}\theta_i, \theta_i)$. Parameterising the Gamma distribution this way so that the shape parameter is clone-dependent (rather than the rate parameter, $v_{ijk} \sim \Gamma(\theta_i, \frac{\theta}{w_{ik}})$), offers closed form distributions and a better fit to the data. The initial number of cells in each well is then sampled from a Poisson distribution $N_{ijk}^{ini} \sim \mathcal{P}(v_{ijk}\lambda_i)$ where $\lambda_i$ is the average total number of cells introduced to wells.

The initial numbers of cells is not observed, so we now model the incubation period. First, for each sample $i$, there are $R_C$ control wells in which we measure the final total number of non-malignant cells $\left(N_{ir}^{C0}\right)_{r\in[R_C]}$ and total number of cells $\left(N_{ir}^{C}\right)_{r\in[R_C]}$. We consider two different survival probabilities for non-malignant and tumour cells, denoted respectively by $\nu_{ir}^{C0}$ and $\nu_{ir}^{C+}$ modelling the drug-independent culture effect. This accounts for various experimental confounding factors, which we detail in the next subsection.

All combined, for the control wells we therefore have

$$\forall r \in [R_C], \quad \left(N_{ir}^{C0}, N_{ir}^{C+}\right) \mid N_{ir}^{C} \sim \text{BetaBin}\left(N_{ir}^{C}, \left(\theta_i w_{i0} \nu_{ir}^{C0}, \theta_i \nu_{ir}^{C+} \sum_{k=1}^{K_i} w_{ik}\right)\right). \tag{8}$$

Finally, for each sample $i$, a drug $d$ is introduced in $R_T$ wells in which we measure the final total number of non-malignant cells $\left(N_{idr}^{T0}\right)_{r\in[R_T]}$ and total number of cells $\left(N_{idr}^{T}\right)_{r\in[R_T]}$. The corresponding survival probabilities for non-malignant and tumour cells are denoted by $\nu_{idr}^{T0}$ and $\nu_{idr}^{T+}$.

Similar to the control wells, the final number of tumour cells and non-malignant cells in wells with drug given the total number of cells is beta-binomial distributed

$$\forall r \in [R_T], \quad \left(N_{idr}^{T0}, N_{idr}^{T+}\right) \mid N_{idr}^{T} \sim \text{BetaBin}\left(N_{idr}^{T}, \left(\theta_i w_{i0} \nu_{idr}^{T0} \pi_{id0}, \theta_i \nu_{idr}^{T+} \sum_{k=1}^{K_i} w_{ik} \pi_{idk}\right)\right). \tag{9}$$

We provide the graphical model corresponding to scClone2DR in Supplementary Figure S17.

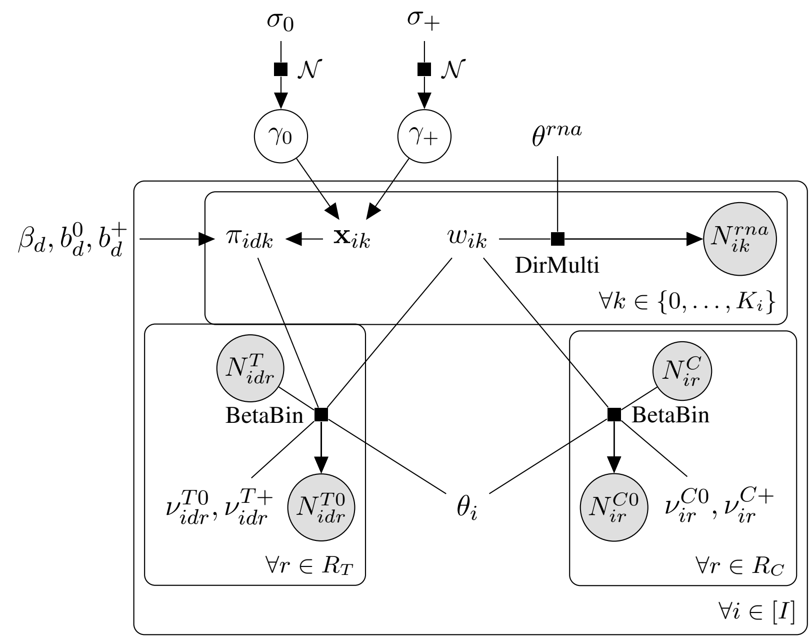


Figure S17: Graphical model of scClone2DR for a specific drug $d$ in the Melanoma cohort, where only two metacell group labels are considered: non-malignant and tumour. The feature vector $\mathbf{x}_{i0}$ (or $\mathbf{x}_{ik}$) for the non-malignant group (or tumour clone $k$) is computed as a learned weighted average (attention mechanism) of the features of all metacells in the corresponding group, parameterized by the latent variable $\gamma_0$ (resp. $\gamma_+$). The survival probability of cells in clone $k$ under drug $d$ is modelled using a sigmoid function applied to the linear predictor defined by the drug-specific regression coefficient $\beta_d$ and the feature vector $\mathbf{x}_{ik}$. Given the latent clone proportion $w_{ik}$ for patient $i$, scClone2DR models the observed cell populations across multiple data modalities of the scRNA measurements, through the clone-specific counts $N_{ik}^{rna}$; and the scFPM measurements, through the total number of cells and the number of non-malignant cells in drug-treated wells ($N_{idr}^{T}$ and $N_{idr}^{T0}$) and in control wells ($N_{ir}^{C}$ and $N_{ir}^{C0}$), across replicates indexed by $r$ on the plate.

### S7 Experimental effects

**Modelling experimental confounding effects is important** We also explore the effect of the experimental design on the scFPM data where cells are placed in wells on a plate, which are filled column by column using a pipette with 12 tips: first, wells in odd columns are filled, then the pipette is recharged and wells in even columns are filled. For patient $i$ and control well replicate $r$, we define the $\nu$-ratio as

$$\frac{N_{ir}^{C+}}{N_{ir}^{C0}} \cdot \frac{N_{i0}^{rna}}{\sum_{k\geq 1} N_{ik}^{rna}}$$

which serves as a proxy derived from observed data of the the quantity $\nu_{ir}^{C+}/\nu_{ir}^{C0}$ of the scClone2DR model. For each replicate $r$, the corresponding well column on the plate is known. We plot the patient-centred $\nu$-ratios against the column position of the well on the plate (Supplementary Figure S18). Patient-centring was performed by subtracting, for each patient, the mean of the ratio computed over all control wells in order to ensures that inter-patient variability does not confound the observed trend with the column of the well.

To quantify the monotonic relationship between column position of the well and the patient-centered $\nu$-ratio, we performed a Spearman rank correlation test. The test revealed a significant correlation between the two variables (p-value of $p = 0.006$), highlighting the need to model the drug-independent culture effect in scClone2DR.

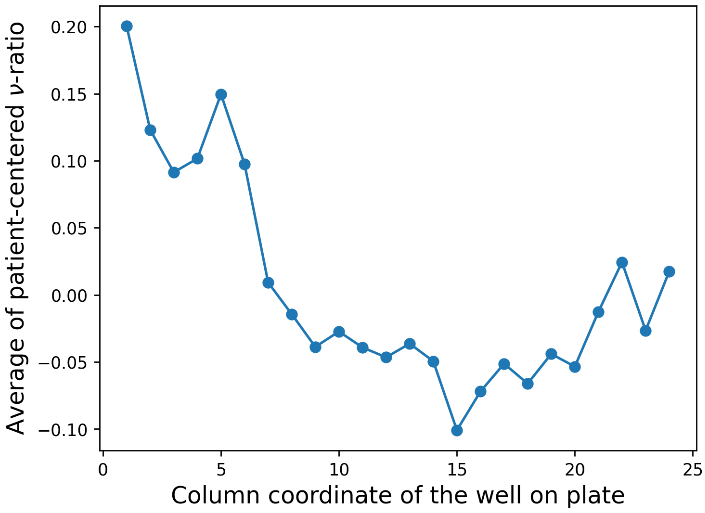


Figure S18: Experimental design introduces a systematic bias in the measured scFPM values: patient-centred $\nu$-ratios depend on the well column of the plate, reflecting the sequence in which wells were filled.

**Modelling and learning the culture effect correction** The drug-independent culture effect is designed to adjust the fraction of non-malignant cells for any factors unrelated to the drug's effect, such as cell density in the well or well position (see Supplementary Figure S18). To model these effects we consider in more detail the process of obtaining the ex-vivo drug screening data:

1. Cells are placed in wells on a plate. Wells are filled column by column. The pipette contains 12 tips. First, wells in odd columns are filled. Then, the pipette is recharged and wells in even columns are filled.
2. Cells stay in suspension then during 24 hours. The position of control wells and the wells in which the drug are added is chosen randomly.

3. After the 24 hours, all cells in all wells are fixed. After fixation, the cells are stained with antibodies that are specifically expressed by cell populations of interest (for example melanoma tumour cells of AML leukemic blasts). Then, the microscope begins capturing images of each well. This is done by scanning from the bottom to the top of one column, then from the top to the bottom of the next column, continuing in this pattern. The entire imaging process takes several hours to complete.

4. Once the images are acquired, cells are classified directly from the fluorescence images (AML) or through a Convolutional Neural Network (melanoma) in order to identify the malignant cells.

The culture survival probabilities $\nu_{idr}^{T0}, \nu_{idr}^{T+}$ and $\nu_{ir}^{C0}, \nu_{ir}^{C+}$ are considered to depend on different factors. First, to mitigate biases resulting from the experimental design, we aim to adjust for variations in cell density within wells and for differences in well positions. Second, changes in tumour content may arise from various interactions between non-malignant and malignant cells in suspension within the wells. We account for this effect indirectly by incorporating the observed fraction of tumour cells from the scRNA-seq data as a feature.

To learn how to correct for the drug-independent culture effect, we consider 4-dimensional feature vectors $\mathbf{z}_{ir}^{C}$ and $\mathbf{z}_{idr}^{T}$, where $\mathbf{z}_{ir}^{C}$ is the feature vector corresponding to the $r$-th control well for the patient $i$. Similarly, $\mathbf{z}_{idr}^{T}$ is the feature vector corresponding to the $r$-th well treated with drug $d$ for the patient $i$. In both cases (control or treated wells), the entries of the feature vectors are $i)$ the final total number of cells in the well divided by maximal total number of cells in a well (observed across all patients), $ii)$ the column of the well (normalized by the total number of 24 columns) if the column is odd and 0 otherwise, $iii)$ the column of the well (normalized by the total number of 24 columns) if the column is even and 0 otherwise, and $iv)$ the observed fraction of tumour cells from the integrated scDNA and scRNA data. We model the logarithm of the ratio of the culture effects using a Linear Generalized Additive Model (GAM) with cubic natural B-splines, employing $n_k = 1$ inner knot with the knot sequence $(0, 0.5, 1)$, namely

$$\log\left(\frac{\nu_{ir}^{C+}}{\nu_{ir}^{C0}}\right) = \mathrm{GAM}(\mathbf{z}_{ir}^{C}) = \eta^\top \mathbf{x}_{ir}^{C}, \qquad \log\left(\frac{\nu_{idr}^{T+}}{\nu_{idr}^{T0}}\right) = \mathrm{GAM}(\mathbf{z}_{idr}^{T}) = \eta^\top \mathbf{x}_{idr}^{T}, \tag{10}$$

where $\mathbf{x}_{ir}^{C}$ and $\mathbf{x}_{idr}^{T}$ are the spline-basis vectors coming out of the GAM model using the feature vectors $\mathbf{z}_{ir}^{C}$ and $\mathbf{z}_{idr}^{T}$.

The penalty matrix $S$ in Equation (6) is correspondingly defined by the following Kronecker product

$$S := \mathbf{Id}_4 \otimes \Omega,$$

where $\mathbf{Id}_4 \in \mathbb{R}^{4\times 4}$ is the identity matrix induced by the four features used in the GAM to model the confounding factors, and where the *per-term penalty matrix* $\Omega \in \mathbb{R}^{(n_k+4)\times(n_k+4)}$ is defined by

$$\Omega_{ij} = \int_0^1 b_i''(t)\, b_j''(t)\, dt = \sum_{u=0}^{n_k} \int_{t_u}^{t_{u+1}} b_i''(t)\, b_j''(t)\, dt, \text{ i.e. } \Omega = 12 \begin{bmatrix} 8 & -11 & 2 & 1 & 0 \\ -11 & 16 & -4 & -2 & 1 \\ 2 & -4 & 4 & -4 & 2 \\ 1 & -2 & -4 & 16 & -11 \\ 0 & 1 & 2 & -11 & 8 \end{bmatrix},$$

with $t_0 = 0, t_1 = 0.5, t_2 = 1$ the knot sequence, and $n_k = 1$ inner knot. Since we use cubic B–splines $(b_i)_i$, $b_i''(t)$ is linear on each interval $[t_u, t_{u+1}]$, hence each interval integral is analytic. The B–spline basis contains $n_k + d + 1 = 1 + 3 + 1 = 5$ elements, where $d = 3$ is the degree of the splines.
